# Evolutionary innovations through gain and loss of genes in the ectomycorrhizal Boletales

**DOI:** 10.1101/2021.09.09.459636

**Authors:** Gang Wu, Shingo Miyauchi, Emmanuelle Morin, Alan Kuo, Elodie Drula, Torda Varga, Annegret Kohler, Bang Feng, Yang Cao, Anna Lipzen, Christopher Daum, Hope Hundley, Jasmyn Pangilinan, Jenifer Johnson, Kerrie Barry, Kurt LaButti, Vivian Ng, Steven Ahrendt, Byoungnam Min, In-Geol Choi, Hongjae Park, Jonathan M. Plett, Jon Magnuson, Joseph W. Spatafora, László G. Nagy, Bernard Henrissat, Igor V. Grigoriev, Zhu-Liang Yang, Jianping Xu, Francis M. Martin

## Abstract

- In this study, we aim to identify genomic traits of the transitions to the ectomycorrhizal ecology within the Boletales, one of the most diverse lineages of symbiotrophic fungi.
- We sequenced the genomes and compared the gene repertoires of symbiotrophic Boletales species to their saprotrophic brown-rot relatives. We also reconstructed gene duplication/loss histories along a time-calibrated phylogeny.
- We showed that the rate of gene duplication is constant along the backbone of Boletales phylogeny with large loss events in lineages leading to several families. The rate of gene family expansion sharply increased in the late Miocene and mostly took place in Boletaceae.
- Most of the ectomycorrhizal Boletales are characterized by a large genome size due to transposable element (TE) expansions and a reduction in the diversity of plant cell wall degrading enzymes (PCWDEs) compared to their brown-rot relatives. However, several species in the Boletaceae, Paxillaceae and Boletinellaceae have kept a substantial set of endoglucanases and LPMOs acting on cellulose/hemicellulose and fungal polysaccharides suggesting that they may partly decompose organic matter by a combined activity of oxidative and hydrolytic enzymes.
- The present study provides novel insights on our understanding of the mechanisms that influence the evolutionary diversification of boletes and symbiosis evolution.

## INTRODUCTION

In forest soils, wood decayers, litter/soil decomposers and ectomycorrhizal fungi are forming entangled networks of mycelia that compete for nutrient resources. A major part of soil carbon and nitrogen organic compounds is stored in forest biomes as wood and soil organic matter (SOM). Decomposition of these resources is thought to be mainly driven by fungal wood decayers and soil/litter saprotrophs (Baldrian *et al.,* 2012). These functional guilds of fungi play a major role in carbon recycling and sequestration, as well as other nutrients, from plant litter and dead bacterial, fungal and animal materials. In addition, ectomycorrhizal symbionts impact SOM decomposition by competing with saprotrophs and regulate the input of plant-related carbon compounds into soil microbial communities. Deciphering how nitrogen acquisition and SOM decomposition potentials differ in saprotrophic and symbiotrophic fungi can provide insight into the underlying mechanisms driving fungal ecological processes and ecosystem functioning (Peay *et al*., 2016). Describing how trait variation and gene copy number of key proteins involved in SOM decomposition and mycorrhizal symbiosis vary across functional guilds will shed light on potential evolutionary trajectories of life history traits in these forest fungi (Lebreton *et al*., 2021).

In the most comprehensive phylogenetic analysis on evolution of ectomycorrhizal fungi in Agaricomycetes carried out to date, 36 origins of ectomycorrhizal lineages were found (Sánchez-García *et al*., 2020). Large-scale studies of mycorrhizal genomes have shown that ancestors of ectomycorrhizal fungi were genetically and ecologically diverse being white rots, brown rots or soil/litter saprotrophs (Kohler *et al*., 2015; Miyauchi *et al*., 2020; Lebreton *et al*., 2021). While tens to hundreds of millions of years separate the independent evolution of ectomycorrhizal symbioses in Endogonales, Ascomycota and Basidiomycota, they share remarkable phenotypic and metabolic similarities. The polyphyletic evolution of the ectomycorrhizal lifestyle is marked by the convergent, near complete loss of core cellulose- and lignin-acting CAZymes, such as ligninolytic peroxidases (class II PODs), cellobiohydrolases of glycosyl hydrolase (GH) families 6 and 7 and cellulose-binding modules found in the ancestral decomposition apparatus of their saprotrophic ancestors. Nevertheless, many of the investigated ectomycorrhizal symbionts have kept a unique array of PCWDEs, including endoglucanases (GH5), pectinases (GH28) and oxidoreductases / laccases (AA1, AA9), thus suggesting that they possess diverse abilities to scavenge plant and microbial detritus from the soil/litter. Finally, TEs and mycorrhiza-induced small secreted proteins (MiSSPs) tend to be enriched in ectomycorrhizal genomes (Pellegrin *et al.,* 2015; Miyauchi *et al*., 2020; Lebreton *et al*., 2021).

With the increasing number of fungal genomes available, it is becoming more feasible to trace the evolutionary events defining the origin of ectomycorrhizal symbiosis within a specific fungal lineage. Functional traits, such as gene copy numbers for PCWDEs, nitrogen and phosphate acquisition pathways, MiSSPs and TEs, within a fungal family across functional guilds (e.g. saprotrophs *vs.* symbiotrophs) have been investigated within the Amanitaceae, Endogonaceae, Russulaceae, Suillaceae, Tuberaceae (Wolfe *et al*., 2012; Murat *et al*., 2018; Chang *et al*., 2019; Loftgren *et al*., 2021; Looney *et al*., 2021). These studies have shown that the abovementioned hallmarks of the ectomycorrhizal ecology are recapitulated at the family level, although idiosyncrasies have been identified in each of these fungal families, e.g. a significant enrichment of terpene- and nonribosomal peptide-synthetase (NRPS)-like secondary metabolism clusters in ectomycorrhizal *Suillus* species.

The order Boletales is among the most species-rich orders in the Agaricomycetes and includes five suborders (i.e., Boletineae, Sclerodermatineae, Suillineae, Coniophorineae and Tapinellineae) and 16 families. Among them, nearly 2/3 of the species are affiliated to the iconic ectomycorrhizal Boletaceae family, as a result of their rapid evolutionary diversification with their co-evolving angiosperm hosts (Grubisha et al., 2001; Binder & Hibbett, 2006; Drehmel et al., 2008; Kirk *et al*., 2008; Dentinger et al., 2010; Sato & Toju, 2019). Species in the suborders Boletineae, Sclerodermatineae and Suillineae establish symbiotic associations with diverse host plants (Newman & Reddell, 1987; Bougher, 1995; Henkel *et al*., 2002; den Bakker *et al*., 2004; Sato *et al*., 2007; Hosen *et al*., 2013; Nuhn *et al*. 2013; Wu *et al*., 2014; Wu *et al*., 2016). On the other hand, species in the suborders Coniophorineae and Tapinellineae are brown-rot fungi that grow on dead wood. No white-rot species are known in Boletales and the most recent common ancestor (MRCA) of Boletales was presumably a brown-rot fungus (Ruiz-Dueñas *et al*., 2020). However, the mode of nutrition in several Boletinellaceae genera, such as *Phlebopus* and *Boletinellus*, is still elusive (Binder & Hibbett, 2006; Tedersoo & Smith, 2013; Sato & Toju, 2019). For example, both the ^13^C/^15^N isotopic signature of the basidiocarps from *P. portentosus* (Berk. & Broome) Boedijn and the ability of its soil mycelium to establish ectomycorrhiza-like structures with *Pinus kesiya* support a symbiotrophic mode of nutrition (Pham *et al*. 2012; Kumla *et al*., 2016). On the other hand, the free-living mycelium of *P. portentosus* is able to produce basidiocarps in the absence of any host tree (e.g., in pots) (Ji *et al*., 2011), suggesting a substantial saprotrophic ability to sustain the massive demand for carbon compounds required by fruiting body construction. These contrasted ecological features suggest that these Boletinellaceae have not fully transitioned to the symbiotrophic lifestyle. Symbiotic Boletales bear several ecologically relevant attributes that warrant study in a genomic context, such as their abundance in temperate, subtropical and tropical ecosystems (Heinemann, 1951; Corner, 1972; Bessette *et al*., 2000; Muñoz, 2005; Zang, 2006), their late-stage fruiting in forest successions (Ortega-Martínez *et al*., 2011), the production of unique volatile organic compounds (VOCs) (Rapior *et al*., 1997) and an accelerated evolutionary rate of speciation, morphological transition and host expansion (Wu *et al*., 2016; Sato & Toju, 2019).

To address a number of hypotheses on the evolution of ectomycorrhizal symbioses from brown-rot ancestors and trophic ecology in the various Boletales families, we sequenced, annotated and compared the genomes of 28 Boletales species, including seven newly sequenced genomes from subtropical and tropical forests. This study had four main objectives. First, to generate a robust phylogenomic framework for the Boletales order. Second, to determine whether the genomes of symbiotrophic Boletales show signatures of the ectomycorrhizal ecology, including large genome size due to TE expansions, reduction in the diversity of PCWDEs and diversification of SSPs that might function in the mycorrhizal symbiosis. Third, to infer the contribution of TE expansion to gene innovation or decay of PCWDEs and SSPs, and fourth, to characterize the gene repertoire and transcript profiling of the Boletinellaceae *Phlebopus portentosus,* having a dual saprotrophic /symbiotrophic lifestyle. By comparing genomes of saprotrophic and symbiotic species, we reveal the genetic basis for their contrasted lignocellulose- and protein-degrading abilities. We also identify major differences in their repertoire of secondary metabolism enzymes and we assess the conservation of symbiotic-related traits in this fungal order.

## MATERIALS & METHODS

### Fungal material, genome sequencing, assembly and annotation

Fungal strains used for genome sequencing are described in Table S1. Genomic DNA was extracted with a modified cetyltrimethylammonium bromide (CTAB) protocol as described in (Kohler *et al*., 2015).

The genome of *Boletus reticuloceps* (M. Zang *et al*.) Q.B. Wang & Y.J. Yao (strain BR01), *Butyriboletus roseoflavus* (Hai B. Li & Hai L. Wei) D. Arora & J.L. Frank (strain LA02), *Lanmaoa asiatica* G. Wu & Zhu L. Yang (strain LA01) and *Chiua virens* (W.F. Chiu) Y.C. Li & Zhu L. Yang (strain LA06) were sequenced for this study using the Pacific Biosciences (PacBio) Sequel platform at Nextomics Biosciences (Wuhan, China). Library construction, genome sequencing, genome assembly and gene annotation were carried out using the company standardized pipeline (Lüli *et al*., 2019). Detailed information is provided in the Supporting Information section. The genome of *Phlebopus* sp. (strain FC_14), *Leucogyrophana mollusca* (Fr.) Pouzar (strain KUC20120723A-06) and *Hygrophoropsis aurantiaca* (Wulfen) Maire (strain ATCC 28755) were sequenced for this study at the Joint Genome Institute (JGI) using the PacBio Sequel or Illumina HiSeq sequencing platforms.

Additional information on methods used for genome sequencing, genome assembly and gene prediction may be found online in the Supporting Information section.

### Organismal phylogeny

We retrieved protein sequences from the Boletales portal at the JGI MycoCosm database (https://mycocosm.jgi.doe.gov/boletales/boletales.info.html) and Nextomics Biosciences (Wuhan, China). A phylogenomic tree was constructed using 28 species of Boletales, two of Atheliales, one of Amylocorticiales and one of Agaricales. Two Polyporales species were chosen as outgroup: *Cristinia sonorae* Nakasone & Gilb and *Polyporus brumalis* (Pers.) Fr. We used 434 highly conserved and generally single-copy protein coding genes that have previously proved useful for higher level phylogenetic analysis of fungi (Beaudet *et al*., 2018; https://github.com/1KFG/Phylogenomics_HMMs, JGI_1086 set), for phylogenomic analyses with the PHYling pipeline (https://github.com/stajichlab/PHYling_unified) with the default settings. Out of the 434 conserved orthologous markers, 430 were identified in our dataset with *hmmsearch* (cutoff=1E-10). The identified protein sequence homologs in each species, for each phylogenetic marker, were aligned with *hmmalign* to the marker profile-HMM. The protein alignments were concatenated into a super-alignment with 430 partitions defined by each gene marker. To select the best-fit amino acid substitution models for each partition, we used ModelTest-NG (v0.1.6, Darriba *et al*., 2019). A maximum likelihood inference for our phylogenomic dataset was achieved with RAxML-NG (v0.9.0, Kozlov *et al*., 2019) using a partitioned analysis and 1000 bootstraps replicates.

### Time-calibrated phylogeny

To calibrate the organismal phylogeny, we used the *mcmctree* method implemented in PAML (v.4.8, Yang, 2007) with the independent-rates clock model, a WAG substitution model and approximate likelihood calculation. For each gene we estimated the substitution rate with *codeml* using the corresponding gene alignment and the following parameters : clock=1, model=2, aaRateFile=wag.dat, getSE=0 and Small_Diff = 1e-7. We then set the time unit to 100 million year (Myr) and applied uniform priors on two fossil calibrations with lower and upper hard bounds. The fossil of a suilloid ectomycorrhizal root tip from the middle Eocene (40–60 Myr ago (Mya)) (LePage *et al*., 1997; Varga *et al*. 2019) was used to calibrate the node containing the suborders Suillineae and a quite relaxed secondary calibration point was set to the Boletales most recent common ancestor (MRCA) using the estimated stem age of Boletales (mean: 218 Mya, 95% HPD: 84–279 Mya) from (Han *et al*., 2018). We also constrained the age of the root to be < 500 Mya.

### Genome synteny and rearrangement analysis

For assessing the genome synteny, we used the 10 largest scaffolds of the genome assemblies with the highest contiguity and completeness, mainly from Boletineae species. We excluded the four Boletalineae genomes having the highest assembly fragmentation, i.e. *Xerocomus badius* (= *Imleria badia*) (Xarba1), *Paxillus ammoniavirescens* (Paxam1), *Paxillus adelphus* (Paxru1) and *Melanogaster broomeianus* (Melbro1; Fig S2). Then, we identified syntenic blocks using a custom script, Synteny Governance Overview (*SynGO*), incorporating the R package *DECIPHER* and *circlize* (Hayat *et al*., 2021; Wright, 2015, Gu *et al*., 2014). We measured the mean TE-to-gene distances with statistical support with the R package *regioneR* (Gel *et al*., 2016). Macrosynteny with combined genomic features among the selected species was visualized and evaluated with a custom script, Visually Integrated Numerous Genres of Omics (*VINGO*), incorporating the R package *karyoploteR* (Looney *et al*., 2021; Gel & Serra, 2017). We assessed and displayed associations between TEs and genes using the R package *vegan* (Oksanen *et al*. 2019) and *PCAtools* (Blighe & Lun, 2020).

### Analysis of gene evolution using COMPARE pipeline

All-vs-all search of whole proteomes of 34 species (Table 1) was performed using mpiBLAST 1.6.0 (Darling *et al*., 2003) with 50 % bidirectional coverage filter and an e-value cutoff of 10^-5^. Proteins were then clustered using the Markov Cluster (MCL) algorithm (van Dongen, 2000) with an inflation parameter 2.0. MCL clusters of protein sequences were aligned using *mafft* 7.4.07 (Katoh & Standley, 2013) with the *--Li-NSI* algorithm and trimmed using *trimAl* 1.4 with a parameter –gt 0.2. Maximum likelihood gene trees for each cluster and Shimodaira-Hasegawa-like (SH-like) branch support values were inferred using the PTHREADS version of RAxML 8.2.12 (Stamatakis, 2014) under the PROTGAMMAWAG model. Next, we reconciled the rooted gene trees with the species tree using Notung 2.9 (Chen *et al*., 2000) with an edge-weight threshold of 0.95. We reconstructed the duplication/loss history of all protein clusters across the species tree using the COMPARE pipeline (Nagy *et al*., 2014; Nagy *et al*., 2016). Duplication and loss rates were computed by dividing the number of inferred duplication and loss events by the length of the respective branch of the species tree. Gene families were functionally characterized by InterPro annotations. We performed InterPro search using InterProScan-5.36-75.0 (Jones *et al*., 2014) with “-goterms” argument. Cluster-based GO enrichment analysis was carried out by using the *topGO* v.2.38.1 package (Alexa & Rahnenfuhrer, 2020) with the *weight01* algorithm (Alexa *et al*., 2006) and Fisher’s exact test. Graphical maps of gene duplication/loss histories were generated using custom scripts in R.

**Table 1.**
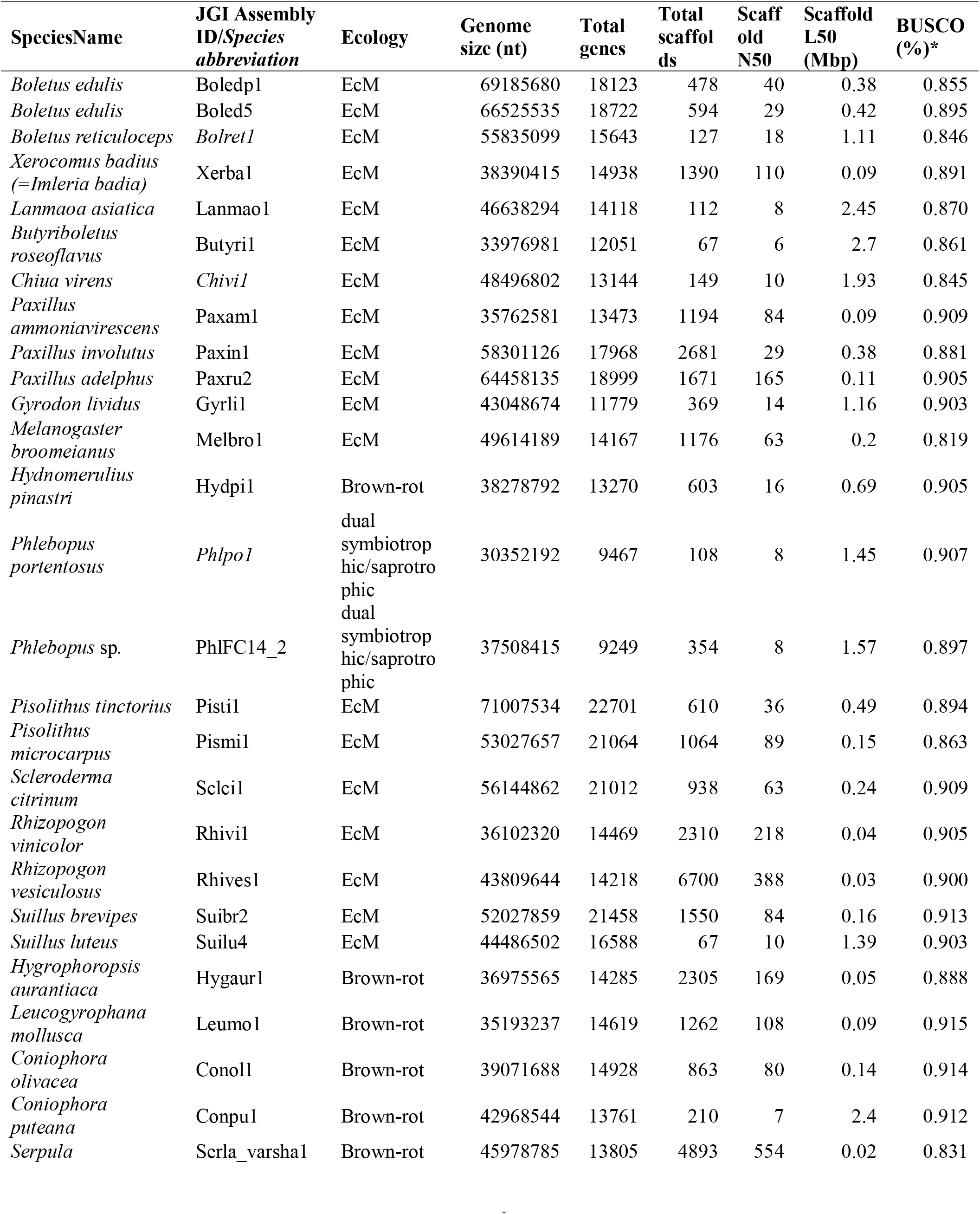
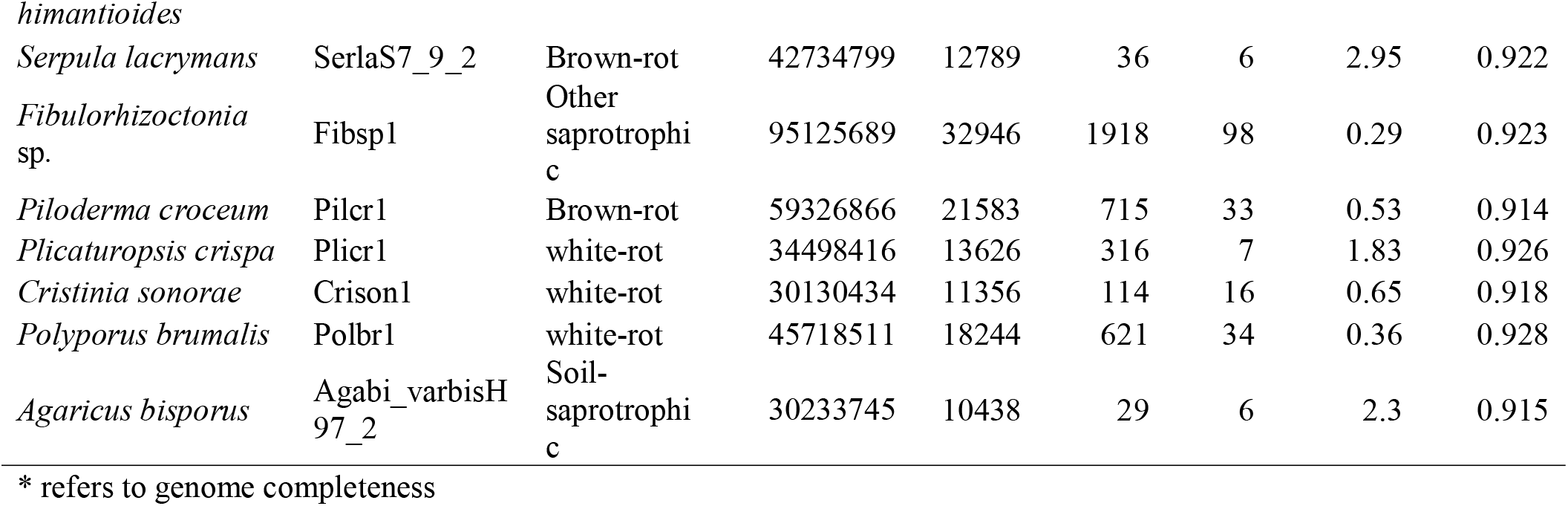
Taxonomic affiliation, brief genomic features and statistics for the 34 included genomes.

## RESULTS

### Genome features and phylogenetic analysis

The nuclear genome of four symbiotrophic Boletaceae (*Boletus reticuloceps*, *Butyriboletus roseoflavus*, *Lanmaoa asiatica* and *Chiua virens*), a dual symbiotrophic/saprotrophic Boletinellaceae (*Phlebopus* sp. FC_14), and two saprotrophic Hygrophoropsidaceae (*Hygrophoropsis aurantiaca* and *Leucogyrophana mollusca*) were newly sequenced, *de novo* assembled and annotated (Table S1) for this study. These genomes were then compared to available sequenced genomes of Boletales in Paxillaceae, Sclerodermatineae, Suillineae, Coniophoraceae and Serpulaceae (Table 1, Table S2). In Boletales, the genome size ranges from 30 Mbp for *Phlebopus portentosus* to 71 Mbp for *Pisolithus tinctorius* (Fig. 1b, Table 1). The completeness of these genome assemblies varies from 83.1%–99.6% in terms of their BUSCO scores with less than 5% missing BUSCO genes (Fig. 1b, Table 1). By combining homolog-based, *ab initio* and transcriptome-based approaches, 9,749 (*P. portentosus*) to 22,701 (*Pi. tinctorius*) protein-coding genes were predicted for these genomes (Fig. 1b, Table 1).

**Figure 1.**
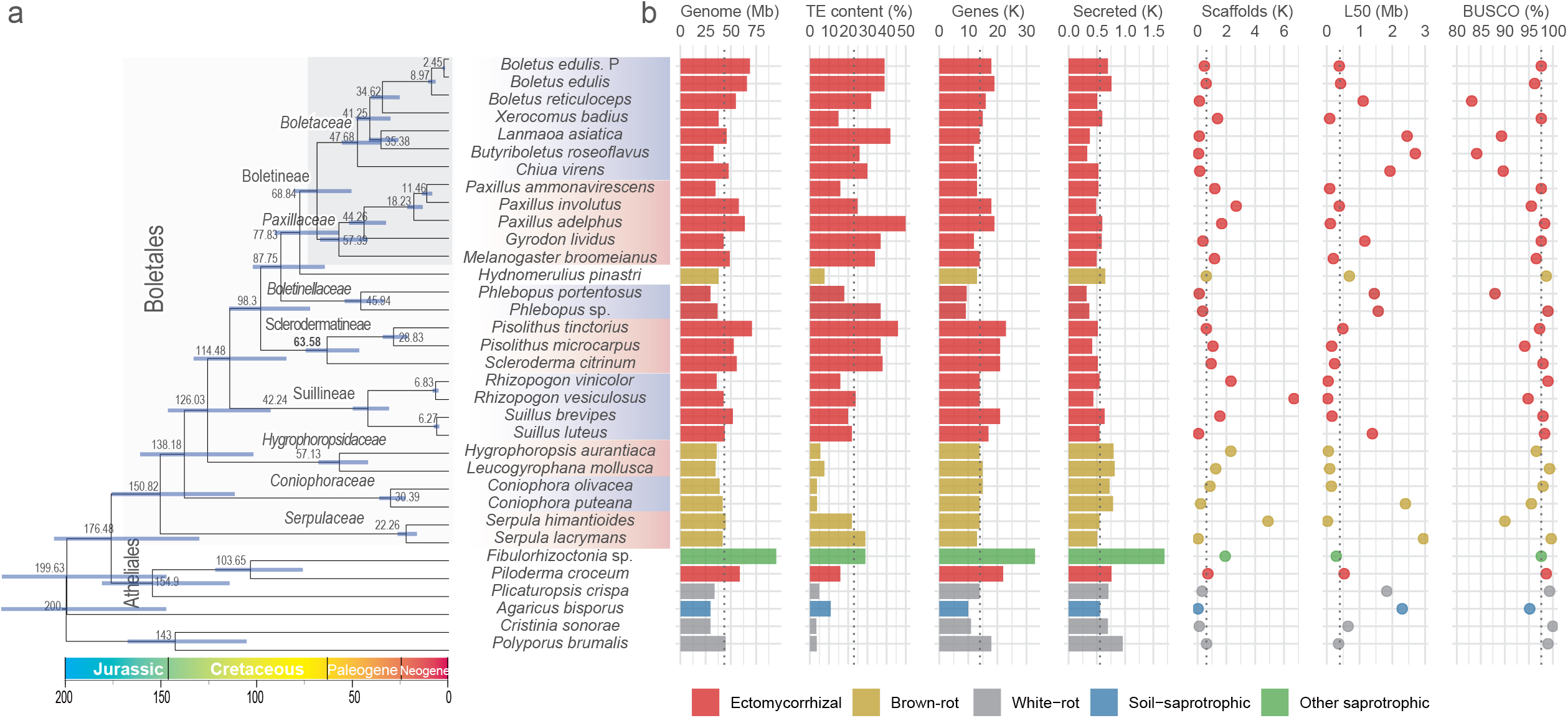
Time-calibrated phylogeny and genome features of 28 Boletales species and six outgroup species. a) The time-calibrated phylogenetic tree in which all nodes received 100% bootstrap support based on maximum likelihood analysis, except the node comprising *Fibulorhizoctonia* sp., *Piloderma croceum*, *Plicaturopsis crispa* with 99% support. The divergence time of each node with confidential interval is shown on/beside the branch. The named clades on the branches of the tree indicate the different taxonomic levels in different fonts (italic font: families, regular font: suborders, regular font in larger size: order). b) Genome features. Colors in plots represent different lifestyles. Genome: Genome size; TE content, coverage of transposable elements in genomes; Genes, number of genes; Secreted, number of secreted proteins; Scaffolds, number of scaffolds; L50, N50 length; BUSCO, BUSCO score indicating genome completeness. The geological timescale (in million year, Myr) is indicated at the bottom.

A Maximum Likelihood phylogenetic analysis, based on the concatenated amino acid sequence alignment of 430 conserved single-copy, orthologous proteins of Boletales and related Atheliales species (Fig. 1a), corroborated and extended previous phylogenetic topologies (Binder *et al*., 2010; Kohler *et al*., 2015; Sato & Toju, 2019). We showed that *H. pinastri,* a brown-rot species, is clustered with ectomycorrhizal Boletineae and Boletinellaceae having a dual symbiotrophic/saprotrophic ecology. The Hygrophoropsidaceae was confirmed as the sister lineage of the main clade comprising Boletineae, Sclerodermatineae and Suillineae (Fig. 1a). The estimated age of the most recent common ancestor (MRCA) of Boletales was 151 (95% PDH: 112–183) Mya in the late Jurassic (Fig. 1a) according to our Bayesian molecular clock dating. The inferred ages were similar to previous estimates (146 Mya, Zhao *et al*., 2017; 142 Mya, Varga *et al*., 2019).

### Genomes of ectomycorrhizal Boletales have a higher load in transposable elements

Ectomycorrhizal fungi have the largest genomes with a significantly higher TE content compared to their brown-rot and soil saprotroph relatives (Fig. 1b; Fig. 2; Tables S3, S4 and S5; p-value < 0.05 for TEs; pair-wise PERMANOVA). TE content ranges from 3.8 % (*Coniophora olivacea*) to 50.3% (*Paxillus adelphus*) of the assembly (Fig. 1b; Table 1, Table S2). The genome size variation was largely explained by the fungal lifestyle (29.3%; Fig. 2; p-value < 0.05; PERMANOVA; Table S4). Among the known TEs, *Gypsy* and *Copia* LTR retrotransposons are widely distributed in Boletales and massively expanded in symbiotic species with lineage-specific features (Fig. 3; Fig. S1, Table S6). Notably, there is an unexpectedly high copy number of *Gypsy*, *Copia* and *EnSpm/CACTA* in *Gyrodon lividus*, an alder-specific symbiont. *Lanmaoa asiatica* genome encodes a very high number of *Copia, Gypsy, hAT* and *Mariner* elements, whereas *C. virens* has a high content in *Copia, Harbinger, Academ* and *Kolobok* families. In contrast to other brown-rot Boletales, *Serpula himantioides* and *S. lacrymans* genomes encode expanded TE repertoires, enriched in *Gypsy* and *Copia* LTR retrotransposons (Fig. 1b, Fig. S1, Table S6). The Kimura distance-based copy divergence showed that TE copies proliferated in the recent historical period in most Boletales species, especially in the ectomycorrhizal symbionts (Fig. 3). Furthermore, a various set of TE families accumulated in very recent time in *Bu. roseoflavus*, *C. virens* and *S. luteus*, while *La. asiatica* and *Pa. involutus* showed two bursts of TE proliferation (Fig. 3). Although TE invasion, proliferation and decay took place at different pace in the various species, it appears that the symbiotic lifestyle is correlated to a higher TE accumulation.

**Figure 2.**
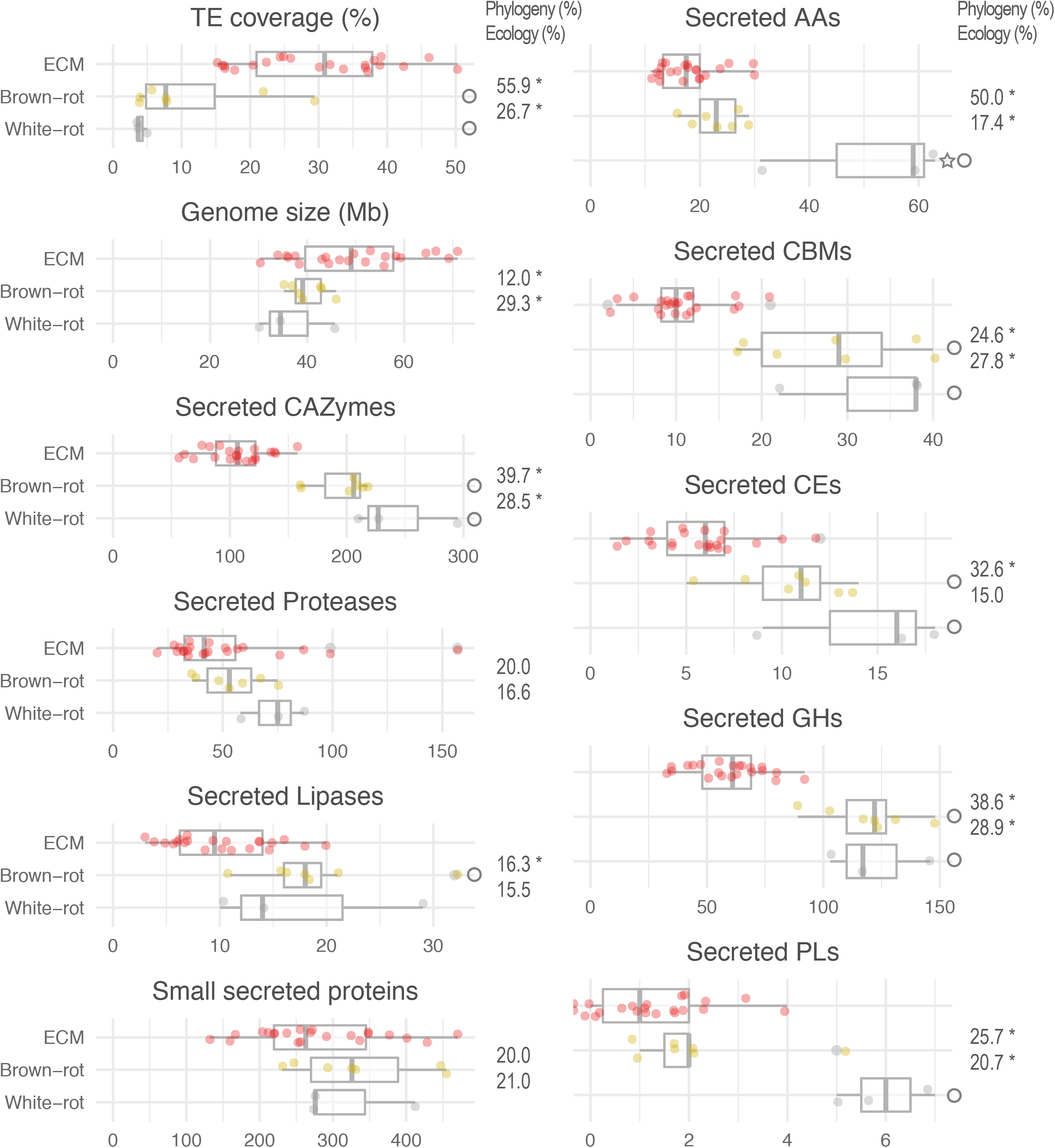
Distribution of the number of secreted proteins with percentage of variance explained by phylogeny and ecological groups. TE coverage, percentage of transposable elements (TEs) in genomes; Genome size, size of genomes in megabase pairs; CAZymes, number of secreted CAZyme domains; Small secreted proteins; number of small secreted proteins (< 300 amino acids); Proteases, number of secreted proteases; CAZyme families (AA, CBM, CE, GH, PL), number of secreted CAZyme domains. Asterisks indicate significantly different phylogenetic distances of species and ecological groups (p-value < 0.05; PERMANOVA model, Genomic features ∼ Phylogeny + Ecology). Circle and star shapes indicate significantly different ecological groups compared with ectomycorrhizal symbionts and brown-rots respectively (p-value < 0.05; Pair-wise PERMANOVA). See detailed information in Table S4 (PERMANOVA; Pair-wise PERMANOVA).

**Figure 3.**
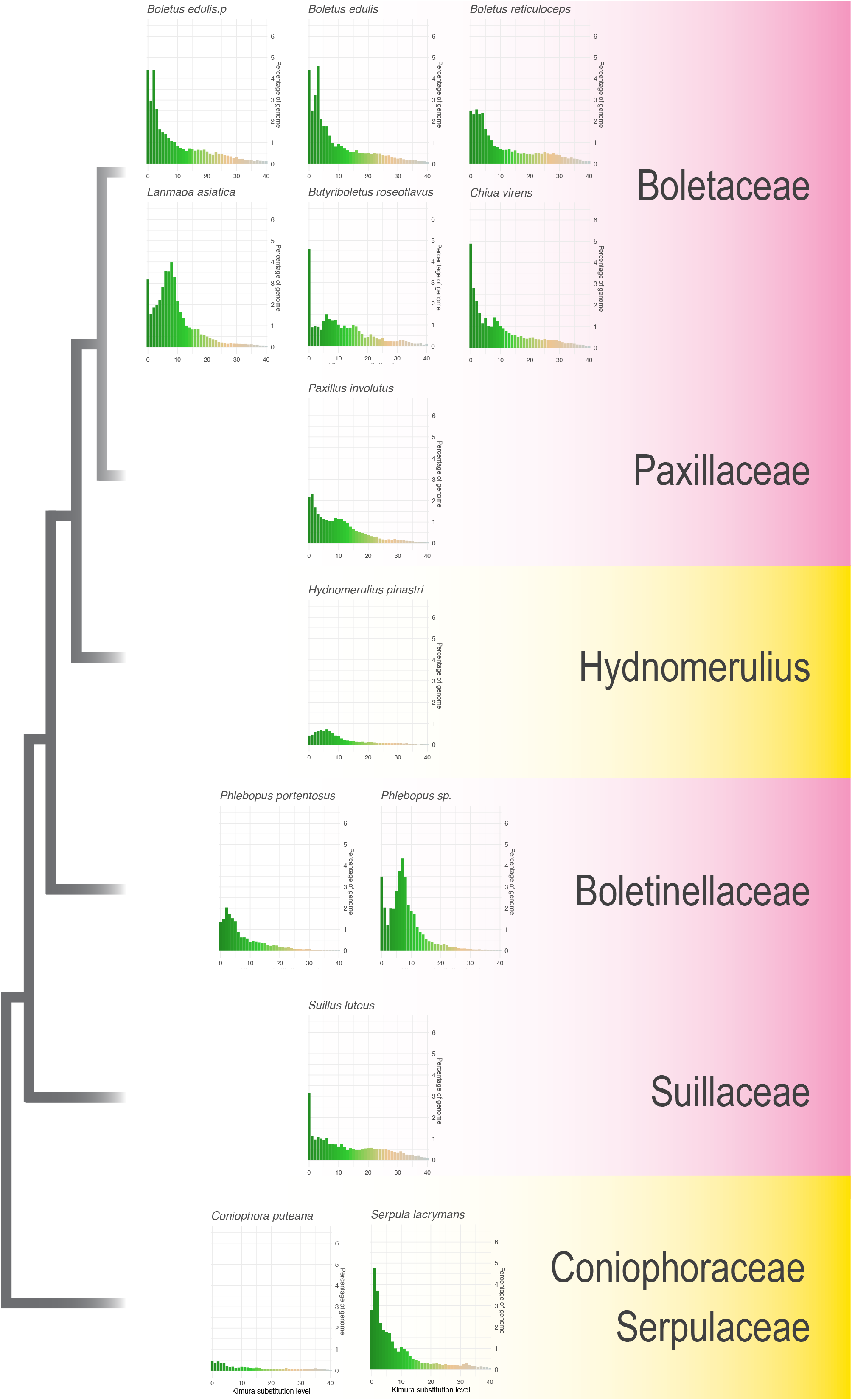
The landscape of transposable elements. (TEs). The bar charts exhibit the genome coverage (%) of all TEs identified on the Y axis. Kimura substitution level (%) on the X axis indicates TE expansion events. Gradient colours (green to grey) show recent to ancient events. The selected representative fungi are sorted in the approximate evolutionary order. The taxonomic family of species are labeled. The fungal lifestyles are in color. Ectomycorrhizal (red), brown-rot (yellow).

### Genomic synteny and rearrangements

We restricted the genome synteny analysis to the top10 largest scaffolds, with the highest contiguity and completeness, including > 50 % of the genome assembly and capturing more than one third of secreted CAZyme coding genes (Fig. S2, Table S7). As expected, the genomes of the European isolates Přilba and BED4 of *B. edulis* share very similar gene repertoires (18722 and 18123 genes, Fig. 1b and 4). Together with the phylogenetically-related *B. reticuloceps*, they present the highest proportion of syntenic regions (Fig. 5ab, Fig. S3, Fig. S4). The genome synteny is however disrupted by numerous TE sequences, suggesting that TE activity reshuffled large parts of these closely-related *Boletus* genomes (Fig. 5ab, Fig. S5, Fig. S6, Fig. S7, Fig S8, Fig S9). The length of syntenic regions decreased with increased phylogenetic distances (Fig. S4). Scaffolds 2 and 5 of *B. edulis* (Boled5) still share many syntenic regions with other Boletales genomes (Fig. S6, Fig. S7, Fig. S9). The CAZyme gene order is well conserved amongst species, but several inversions took place (e.g., between *Pa. involutus* (Paxin1), *H*. *pinastri* (Hydpi1) and *P. portentosus* (Phlpo1)). Intriguingly, gene insertions or inversions were often observed in TE-rich regions (e.g., the genes coding for AA3_2, EXPN, GH128 and GT2 labelled in orange (Fig. S6, Fig. S7, Fig. S8), suggesting that TE insertions played a key role in these events.

**Figure 4.**
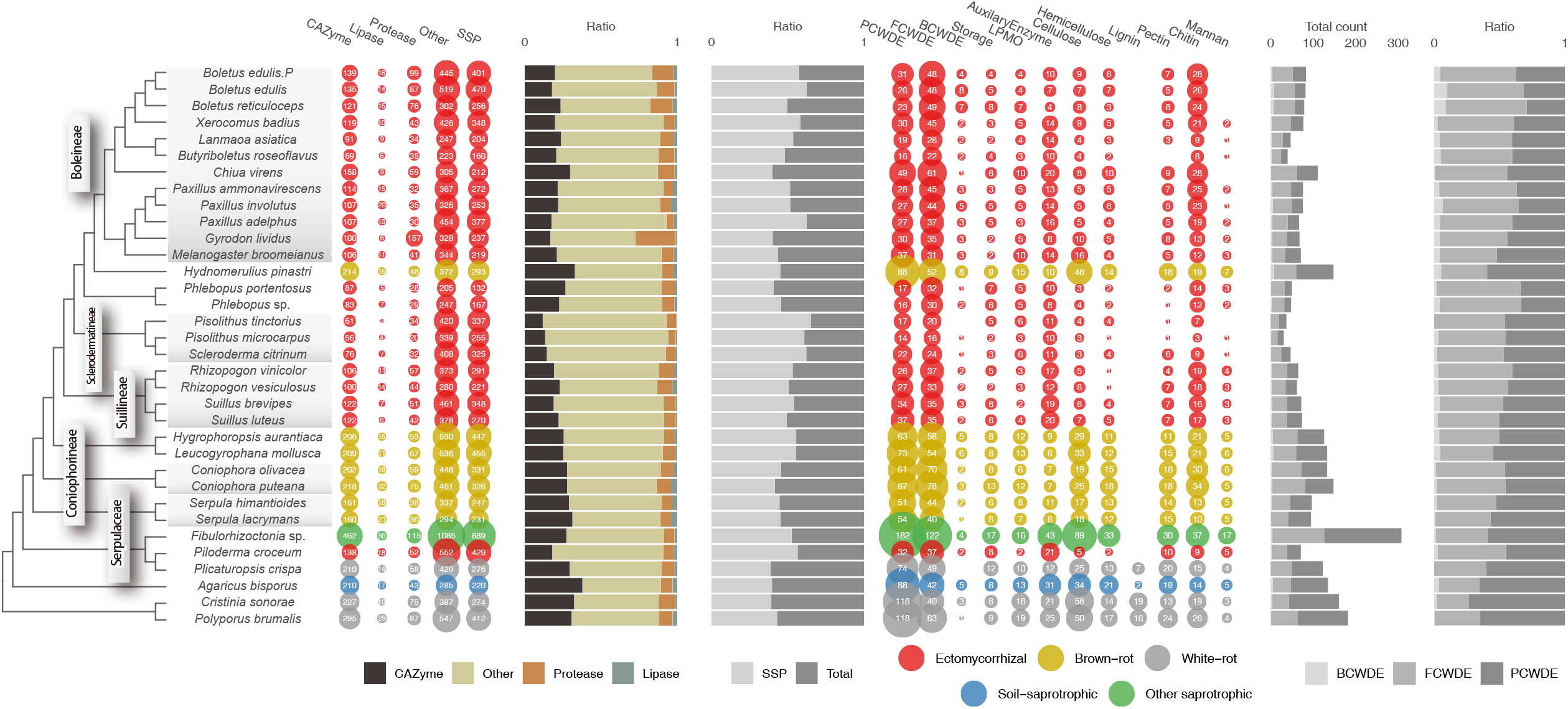
Secretome profiles of 28 Boletales and six outgroup species. The first bubble plot (on the left) shows the number of secreted genes for CAZymes, lipases, proteases and others (i.e. all secreted proteins not in these first three groups). The SSP group is a subcategory showing the number of SSPs (< 300 aa). The size of the bubbles corresponds to the number of genes. Fungal species are color coded according to their known ecology. The first bar plots (in the middle) represent the ratio of CAZymes, lipases, proteases, to all secreted proteins (left) and the ratio of SSPs among the entire secretome (right). The second bubble plot (on the right) shows the number of plant cell wall degrading enzymes (PCWDE) and microbial cell wall degrading enzymes (MCWDE, including bacterial cell wall degrading enzymes (BCWDE) and fungal cell wall degrading enzymes (FCWDE)), lytic polysaccharide monooxygenases (LPMO), substrate-specific enzymes for cellulose, hemicellulose, lignin, and pectin (plant cell walls); chitin, glucan, mannan (fungal cell walls). The second bar plots (far right) show the total count of genes including PCWDE and MCWDE (left) and the ratios of PCWDE, BCWDE and FCWDE (right).

Of note, we found that genes coding for proteases, lipases and SSPs are closer to the nearest TEs than the random expectation (Fig. 5c, Table S8). Multivariate and ordination analyses further indicated that unclassified TE sequences are significantly associated with gene counts for chitin synthase (GT2) and SSP genes in ectomycorrhizal genomes (Fig. S10, Table 2, Table S9).

**Figure 5.**
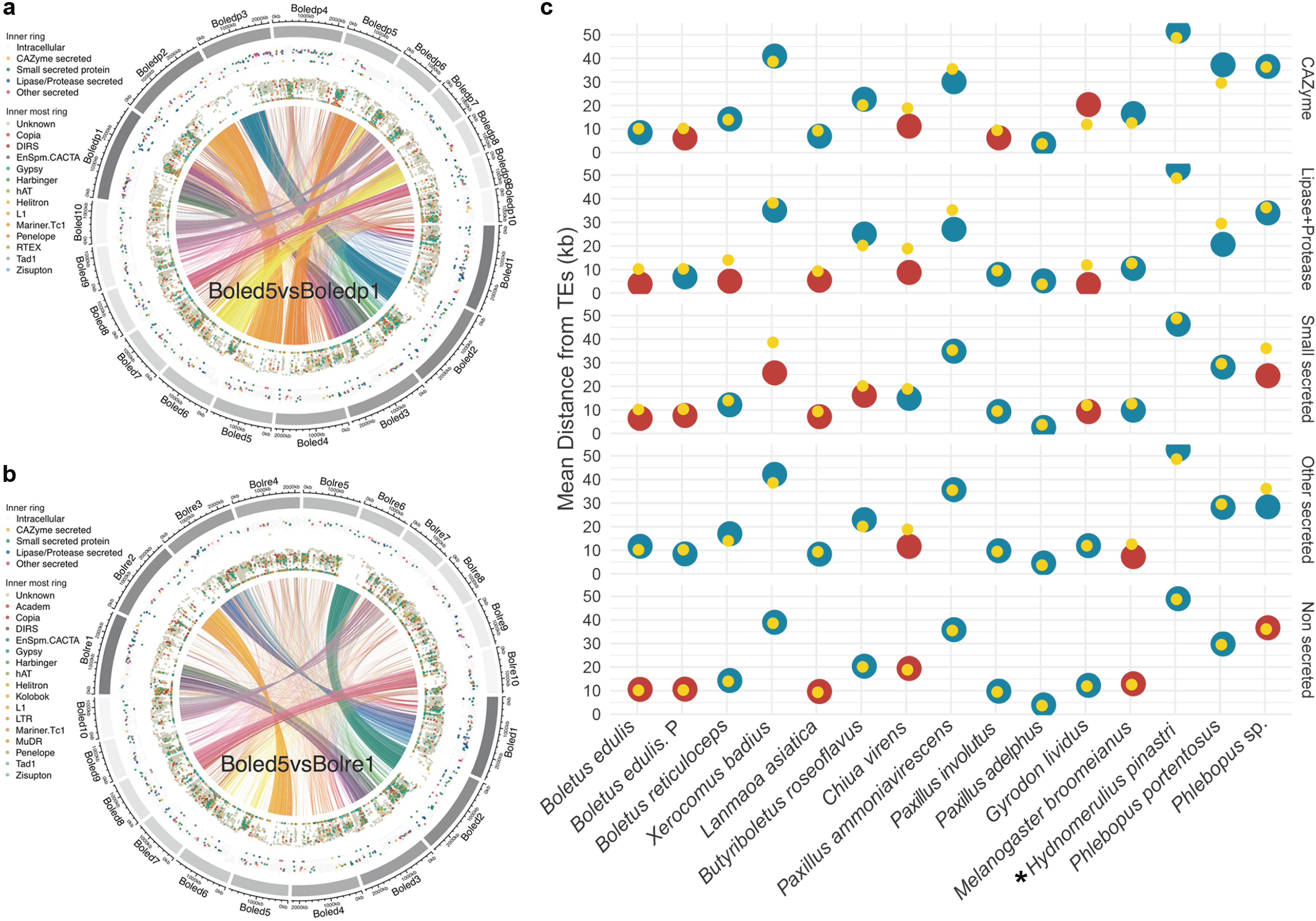
Syntenic analysis of *Boletus* species (a, b) and genomic distance investigation between genes and TEs (c). a) and b) Plots showing genes and TEs present only in syntenic regions of *Boletus* species Outer circle, scaffold size; first inner circle, genes coding for secreted CAZymes, SSPs, lipases, or proteases (see legend); second inner circle, TEs (see legend); vertical axis of inner circles, mean distances of between neighboring genes and TEs. Shorter distances between genes and TEs result in dots towards plot centers, whereas longer distances result in dots towards the outer circle. Boled5, *Boletus edulis* (BED1); Boledp1, *Boletus edulis* P (Přilba); Bolre1, *Boletus reticuloceps* (BR01). c) showing mean distances between TEs and genes coding for CAZymes, lipases/proteases, small secreted proteins, other secreted proteins and non-secreted proteins. Yellow: mean distances of 10,000 randomly reshuffled genome models (to generate a null hypothesis). Blue: mean distances observed in genomes with no statistical significance (p > 0.05). Red: mean distances observed in genomes with statistical significance (p < 0.05). Brown-rot (asterisk), ectomycorrhizal (unmarked). See Supporting Information-Table S8 for detailed mean distances between repeat elements and genes.

**Table 2.** Multivariate analysisa of TEs and genes.

| Variation explained by<br>unclassified repeats (%) | FDR adjusted p values | Count of genes |
| --- | --- | --- |
| 21.90% | p < 0.01 | GT2 |
| 11.70% | p < 0.05 | Total GTs |
| 8.75% | p < 0.05 | SSPs |

<sup>a</sup> See details of PERMANOVA results in Table S9.

A hallmark of the genome evolution in ectomycorrhizal species is the dramatic loss of genes encoding CAZymes acting on plant cell wall polysaccharides in most species. We thus investigate the microsynteny of several CAZyme genes to track down the underlying mechanism(s) leading to gene loss. We found that the conservation of the nucleotide sequences of protein-coding genes framing the CAZyme genes, such as the cellobiohydrolase GH6 (Fig. S11), but also cellobiohydrolase (GH7), endoglucanases (CBM1-GH5_5) and xyloglucanase (GH74-CBM1) (data not shown), is highly conserved with no apparent disruption by TE sequences. It appears that the loss of these genes in ectomycorrhizal Boletales involved the accumulation of discrete nucleotide mutations, the so-called DNA decay process also observed in Tuberaceae (Murat *et al*., 2018).

### Core, dispensable and species-specific gene families in Boletales

To compare the protein-coding gene repertoires encoded by the sequenced Boletales and identify species-specific gene families that might contribute to diversification of symbiosis-related traits in ectomycorrhizal Boletales, we clustered the predicted proteins to infer orthologous gene groups (orthogroups), including core genes (i.e., occurring in the 28 Boletales species), dispensable genes (i.e., found in at least two species) and species-specific genes (i.e., unique to a taxon). While the core set of conserved genes ranged from 2,291 to 2,757, the repertoire of species-specific genes varies widely, ranging from 1,364 in *Phlebopus* sp. FC_14 to 12,816 in *Pi. tinctorius* (Fig. S12a, Table S10). The proportion of species-specific genes is much higher in Sclerodermatineae in comparison to Boletineae. Species-specific genes, which are also referred to as taxonomically-restricted genes have recently become associated with the evolution of novelty, as numerous studies across the tree of life have now linked expression of taxonomically-restricted genes with novel phenotypes (Johnson, 2018). As expected, they mostly encode proteins with no known function in the Boletales. We further identified the orthogroups allowing the discrimination between saprotrophs and ectomycorrhizal symbionts. The number of symbiont-specific orthogroups ranges from 1347 in *G. lividus* to 5231 in *B. edulis* BED1, and that of brown-rot-specific genes varies from 842 in *H. pinastri* to 3755 in *Coniophora puteana* (Fig. S12b, Table S11). The number of symbiont/brown-rot-specific orthogroups tended to be proportional to the genome size of the corresponding fungi (Fig. 1, Fig. S12b).

### Diversification of the gene repertoires

The newly sequenced genomes, and more specifically the Boletaceae and Boletinellaceae genomes, offer a unique opportunity to examine the evolution of bolete-related and symbiosis-related genes, and gene families across the Boletales order. Our reconstructions of genome-wide duplication and contraction events in Boletales revealed a considerable heterogeneity in the temporal dynamics of genome diversification between the different clades (Fig. 6a). We reconstructed similar ancestral copy numbers (12000 to 13000 genes) in ancestral nodes, with a moderate increase from the root of the tree (Fig. 6a). In accordance, net gene duplications (duplications minus losses) were inferred to be more or less constant along the tree backbone with large loss events (2000 to 5000 gene losses, Fig. 6a) on the branches leading to the families. The mean gene duplication rate in Boletales increased abruptly around 10 Mya in the late Miocene (Fig. 6b). This seems to be driven mostly by the family Boletaceae and to a smaller extent by Suillaceae (Fig. S13). This may be explained by at least two hypotheses. First, constraint on gene duplicability may have been lifted in these clades, leading to a surge of gene duplications. Second, single events with large impacts (e.g. large segmental duplications) may have shaped Boletaceae (and Suillaceae) genomes, leading to an increase in duplication rates. Third, it is also possible that the observed gene duplications represent adaptive events that contributed to the evolutionary success of boletes. Which, or whether a mixture, of these can best explain the observed rate increase will need to be examined with more in-depth analyses. It should also be noted that gene duplication/loss rates may be underestimated in our analyses due to the inherent inability of all comparative approaches to account for events in extinct lineages. However, it is unlikely that this has an effect on the detection rate acceleration in the Boletaceae.

**Figure 6.**
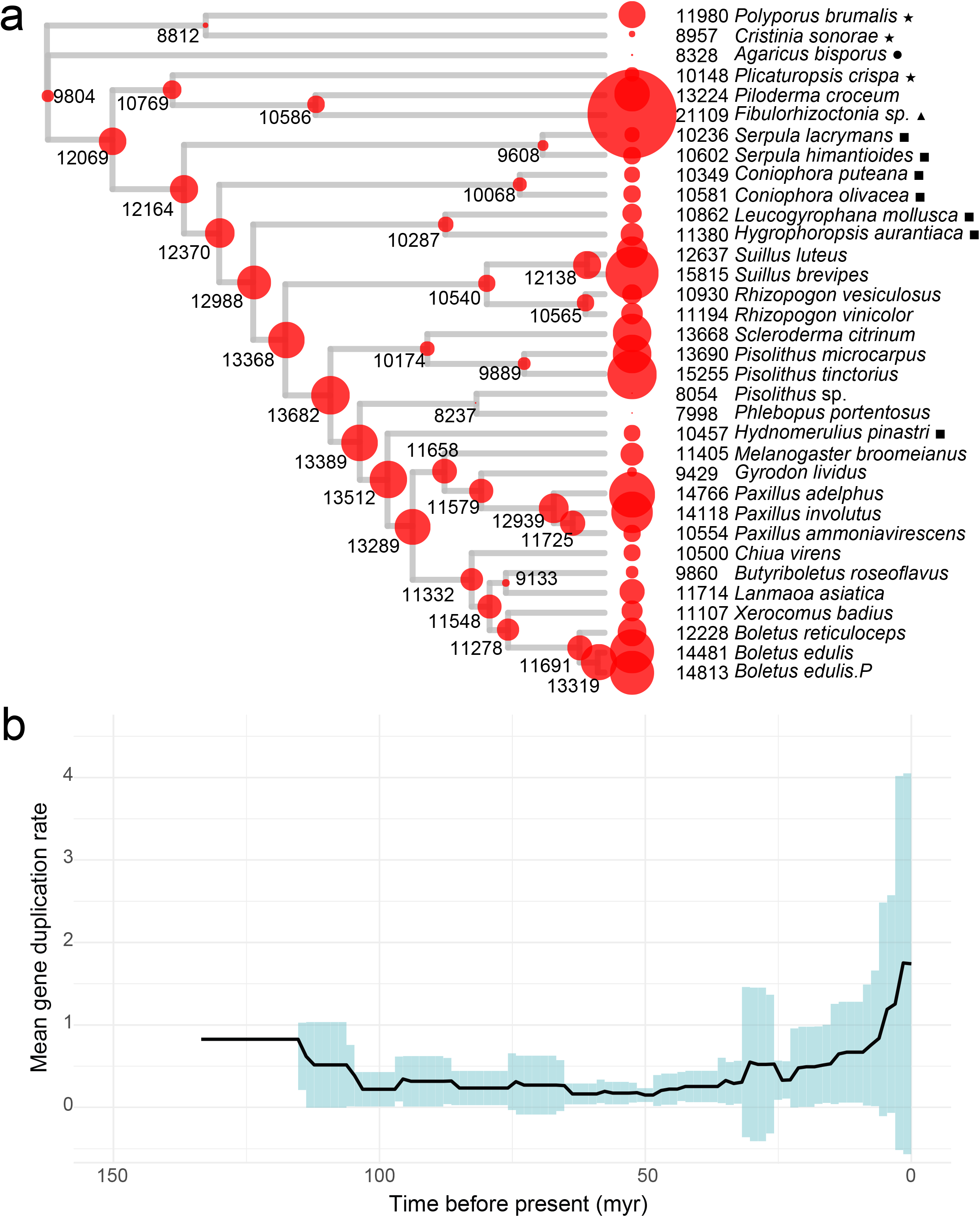
COMPARE analysis of Boletales. **a)** The reconstructed gene copy numbers of Boletales species and outgroup species. **b)** Gene duplication rates through time plot. Solid black line represents the mean of the gene duplication rate across lineages and time. Blue bars depict the standard deviation of the gene duplication rates in a given time frame. Time frames were calculated by dividing the time interval of Boletales evolution by 100. White-rot (asterisk), soil-saprotrophic (circle), brown-rot (quadrangle), other-saprotrophic (triangle), ectomycorrhizal (unmarked).

To gain insights into the unique features of the bolete genomes, we compared GO annotation frequencies of genes having duplications at ancestral nodes of Boletaceae. We performed gene family-based enrichment analysis using GO annotations and found 23 molecular function, 13 biological process and seven cellular component terms, which were significantly overrepresented (Fisher’s exact test, p<0.05) in gene families that showed duplications in early Boletaceae (Table S12). The most significant terms were “superoxide dismutase activity”, “ubiquitin-like modifier activating enzyme activity”, “protein kinase activity “, “proteolysis”, “organophosphate catabolic process”, “myosin complex” and “integral component of plasma membrane”. We found 169 InterPro terms that were significantly enriched (hypergeometric test, p < 0.05) among gene families which showed duplications in ancestors of the Boletaceae. Among these InterPro terms some exhibited specific functions such as those related to protein kinases (e.g., fungal-type protein kinase, serine/threonine-protein kinase), GHs (LPMOs, GH20 β-hexosaminidase and GH18 chitinase) and transposases (*Kyakuja-Dileera-Zisupton* transposase, *Plavaka* transposase and *Tc1*-like transposase).

### Lignocellulose and protein degradation abilities in Boletales

The repertoires of predicted secreted proteins in Boletales species were compared to identify possible lineage-specific features of the machinery involved in SOM and plant/microbial cell wall degradation (Fig. 4, Fig. S14). Ectomycorrhizal Boletales have a significantly smaller mean number of secreted CAZymes (e.g. GHs) compared to brown-rot and white-rot species, while no significant difference was found for SSPs (Fig. 2, Table S4; p-value < 0.05; Pair-wise PERMANOVA). We sorted gene families coding for secreted cell wall degrading enzymes into three main functional categories, i.e. enzymes involved in the degradation of lignin, cellulose/hemicellulose (including pectin) and microbial polysaccharides (i.e., chitin, fungal and bacterial glucans). The PCWDE gene copy numbers showed striking differences between brown-rot and ectomycorrhizal species (Fig. 4). The number of secreted PCWDEs in ectomycorrhizal species ranged from 16 for *Bu. roseoflavus* to 49 for *C. virens*. The non-metric multidimensional scaling (NMDS) analysis grouped the sequenced Boletales species according to their known nutritional modes, brown-rot *vs.* symbiosis (Fig. S15). Phylogenetic relatedness of the species (39.7 %) and fungal ecology (28.5 %) significantly accounted for the variation in secreted CAZyme genes (Fig. 2, Table S4; p-value < 0.05; PERMANOVA model, Genomic features ∼ Phylogeny + Ecology). The ectomycorrhizal Sclerodermatineae (*Pi. tinctorius*, *Pi. microcarpus*, and *Sc. citrinum*) were further separated from other symbiotic species because of their even lower content in secreted CAZymes (Fig. 4, Fig. S15).

Brown-rot and ectomycorrhizal Boletales and Atheliales lack class II lignin-modifying PODs (AA2) (Fig. S14). In addition, the loss of synergistically acting GH7 and GH6 cellobiohydrolases, operating on cellulose reducing and non-reducing ends, is a hallmark of the ectomycorrhizal Boletales (Fig. S14). The near complete loss of core cellulose-acting CAZymes is underlined by the restricted set of CBM1 binding modules, which are often attached to key PCWDEs to mediate the targeting of enzymes to cellulose. However, a single copy of the endoglucanase of families GH9 and GH45, digesting cellulose into cellooligosaccharides, are found in the symbiotrophic species, suggesting a limited capacity to degrade cellulose. The endoglucanase GH12 is missing from *Paxillus*, *Phlebopus* and *Scleroderma* species, but are present in the Boletaceae. Concerning hemicellulose degradation, neither endo-β-1,4-xylanases (GH11) acting on the xyloglucan backbone nor the debranching enzymes working synergistically with GH11 (families GH54, GH62, GH67 and CE3) were found. Enzymes involved in lignocellulose oxidation (LOX) in brown-rot species, such as cellobiodehydrogenases (AA3_1), iron reductase domain containing proteins (AA8), and LPMOs (AA16), are also missing from the symbiotic Boletales genomes, except for one copy of pyranose-2-oxidase (AA3_4) in *B. edulis*. In contrast, gene copy number in several LOX gene families, including laccases (AA1_1), aryl alcohol/glucose oxidases (AA3_2), alcohol oxidases (AA3_3) (except for *Pisolithus* species), glyoxal oxidases (AA5), benzoquinone reductases (AA6), glucooligosaccharide oxidases (AA7), cellulose-acting LPMOs (AA9) and xylan-acting LPMOs (AA14) is similar in brown-rot and symbiotic species. *Chiua virens*, a basal species of Boletaceae, has a very high number of phenoloxidases (AA1_1), even higher than that of Boletales brown-rot species. Similarly, the copy number of genes coding for enzymes acting on fungal and bacterial polysaccharides is similar for ectomycorrhizal and brown-rot Boletales (Fig. 4).

The number of genes coding for secreted proteases ranged from 36 to 75 in brown-rot species, while it varies from 20 to 157 in ectomycorrhizal symbionts (Fig. 4), suggesting they have a similar capacity to degrade proteins accumulating in SOM. However, the set of secreted proteases is much higher in *Boletus* species (*B. edulis*, 87 and 99 copies; *B. reticuloceps*, 76 copies) and *G. lividus* (157 copies) (Fig. 4). We observed a striking gene duplication of the gene encoding pepstatin-insensitive carboxyl proteinases (family G01, formerly A4) in the *Boletus* clade with 25 to 34 copies (Fig. S16). This gene duplication occurred in a syntenic region in *B. reticuloceps* (Bolret1) and *B. edulis* Přilba (Boledp1) (Fig. S8, Table S13). Gene copy number for secreted lipases is similar in the genomes of ectomycorrhizal and brown-rot Boletales (Fig. 4), except for Sclerodermatineae with the lower repertoire of lipases.

### *Phlebopus portentosus* saprotrophic capability

As above mentioned, *P. portentosus* PP33 and *Phlebopus sp.* FC_14 are amongst the ectomycorrhizal Boletales having the lower set of PCWDEs (Fig. 4), suggesting a limited saprotrophic ability. Still, *P. portentosus* PP33 is able to produce basidiocarps in the absence of any host plant (Pham *et al*. 2012; Kumla *et al*., 2016), raising the question of the catabolic pathways used to provide the high load of simple carbohydrates required for sustaining basidiocarp development. We thus performed RNA-seq profiling of *P. portentosus* PP33 free-mycelium grown either on PDA medium or the rubber tree sawdust organic medium used to trigger basidiocarp development and measured the expression of the few genes coding for secreted PCWDEs (Table S14). We found that genes coding for laccase (AA1_1), xyloglucan:xyloglucosyltransferase (GH16), chitinase (GH18), chitin deacetylase (CE4), β-1,3-glucanase (GH128), exo-β-1,3-glucanase (GH55) and cellulose-acting LPMO (AA9) are expressed at a high level on PDA and sawdust/rice seed organic media (Fig. S17, Fig. S18), suggesting the corresponding enzymes are involved in the release of soluble carbohydrates from plant organic matter. The higher expression of the laccase and chitinase genes suggests that sawdust polyphenols and chitin from fungal walls are potentially used to sustain the increased carbon demand required by fruiting body development.

### Conservation of symbiosis-induced genes

To test the conservation of ectomycorrhiza-related genes within Boletales species, we characterized the protein sequence similarity of available symbiosis-induced genes from ectomycorrhizal Boletales species amongst the nine saprotrophic and 19 symbiotrophic Boletales genomes. In the absence of transcriptomic datasets from bolete ectomycorrhizas, we queried the protein-coding gene repertoires using BLASTP and 275 known symbiosis-induced genes retrieved from *Suillus luteus*, *Pa. involutus, Pi. microcarpus* and *Pi. tinctorius* transcript profiles obtained from ectomycorrhizal root tips of *Betula*, *Eucalyptus* and *Pinus* (Kohler *et al*., 2015) (Fig. S19). Most symbiosis-induced genes (85 %) are conserved amongst saprotrophic and symbiotic Boletales (clusters II to V). These conserved symbiosis-induced genes encode for proteins involved in cellular processes and signaling, core metabolic pathways and CAZymes, or proteins with no KOG annotation. The 80 genes (10 %) from cluster I are mainly specific to *Paxillus, Pisolithus* or *Suillus* species. They mainly encode MiSSPs or genes with no known function (no KOG domain).

### Transporters, transcription factors and secondary metabolism enzymes

Although the genomes of the investigated Boletales species have a similar set of terpene cyclases, symbiotrophic boletes have a smaller number of genes encoding non-ribosomal peptide synthetases (NRPS) and NRPS-like enzymes, than brown-rot species (Fig. S20), indicating a restricted capability to synthesize secondary metabolites. The gene copy numbers for the different membrane transporters and transcriptional regulators revealed no specific pattern(s) for ectomycorrhizal fungi by comparison to brown-rot species (Fig. S20).

## DISCUSSION

The Boletales order includes ∼1300 described species (Kirk *et al*., 2008). As either brown-rot decomposers or ectomycorrhizal symbionts, they play a key role in carbon cycling and sequestration in boreal, temperate and mountane forests (Eastwood *et al*., 2011). Understanding the evolution and ecology of this major fungal group can be achieved only if coupled with a comprehensive description of their genomic traits.

Deciphering the evolution of gene families related to the modes of nutrition, i.e., saprotrophy or symbiotrophy, is of particular interest to understand the ecology of these plant-associated fungi. The number of published Boletales genomes is now large enough to permit a comparative analysis of genome evolution between and within suborders and families exhibiting a wide range of ecological traits (e.g., mode of nutrition, habitat, host specificity). The 28 genomes of Boletales compared in the present study cover both the saprotrophic and symbiotic lifestyles found in this order. The evolutionary relationships between the selected Boletales and related Atheliales species were assessed by constructing a time-calibrated phylogenetic tree. Our phylogeny is in agreement with the recent megaphylogenies of Agaricomycetes (Krah *et al*., 2018; Varga *et al*., 2019). In these phylogenies, the emergence of the MRCA of Agaricomycetidae, Agaricales, Polyporales, Russulales and Boletales was dated at 185, 173, 150, 152 and 142 Myr, respectively. In our analysis, the MRCA of the Boletales was dated at 151 Myr in late Jurassic, which is close to previous estimates. Many of the orders within Agaricomycetes (Agaricales, Boletales, Polyporales) originated before, but diversified after, the angiosperm and gymnosperm origins (Krah *et al*., 2018). Hibbett & Matheny (2009) suggested that the ectomycorrhizal Boletales are younger than angiosperms and conifers, but slightly older than the rosids, which contain many symbiotic partners of extant Boletales.

In our analysis, most brown-rot producing Boletales species are placed as a paraphyletic group at the base of Boletales. We however confirmed the placement of *H. pinastri* among ectomycorrhizal species in the “core Boletales” (Sclerodermatineae, Suillineae, and Boletineae) (Kohler *et al*., 2015; Krah *et al*., 2018). This brown-rot species retains numerous PCWDE genes loss in the lineages leading to the ectomycorrhizal members of the “core Boletales” (this study, Kohler *et al*. (2015)), suggesting that the ancestor of the core Boletales had substantial saprotrophic ability, which has been retained in *H. pinastri*. Our work reveals that the overall pattern of gene loss and diversification differs amongst the ectomycorrhizal Boletales suborders, supporting the contention that these lineages originated from different ecologically diverse brown-rot precursors.

In their analysis of the Agaricomycetes evolution, Sánchez-García *et al*. (2020) showed that across their 8,400-species phylogeny, diversification rates of ectomycorrhizal lineages were no greater than those of saprotrophic lineages. However, some ectomycorrhizal lineages have elevated diversification rates by comparison to their nonsymbiotic sister clades, suggesting that the evolution of ectomycorrhizal symbioses may act as a key innovation at local phylogenetic scales. In the Boletales, the relative time course of genome innovation in each of the family-level clades is distinct. PCWDE and SM gene losses is a well-known pattern associated with ectomycorrhizal evolution (Wolfe *et al*., 2012; Martin *et al*., 2016; Hess *et al*., 2018; Lebreton *et al*., 2021). Here, we showed that extensive gene loss took place in the early ectomycorrhizal Boletales clades. A substantial proportion of protein orthogroups contracted within the MRCA of ectomycorrhizal Boletales and appeared purged from the genome completely, suggesting that the early loss of a wide range of genes accompanied the formation of ectomycorrhizal associations. However, the present analysis also suggests a gradual increase in gene duplication rates in Boletaceae that is especially apparent through the late Miocene to present. During this period, TEs also proliferated in most Boletales genomes, although Boletaceae genomes experienced the highest accumulation. Of note, three protein domains related to transposases (IPR040521, IPR038717, IPR041078) are enriched among ancestral nodes of Boletaceae. Our findings support the view that most Boletales species are undergoing a period of genome expansion (Castanera *et al*. 2017). Intriguingly, TE-mediated genome amplification coincides with the estimated origins of ectomycorrhizal symbiosis in Boletales (this study, Kohler *et al*., 2015).

The higher copy number of chitin synthase (*GT2*) genes in Boletaceae could be explained by the activity of TEs as suggested by the close location of several *GT2* genes to TEs. In addition, the genomic location of genes coding for SSPs, proteases and lipases tended to be closer to the nearest TEs than the random expectation. This suggests that TEs have contributed to the observed higher evolutionary rate of genes encoding effector-like SSPs, proteases, and lipases. These genome innovations may be related to the dramatic increase in species-richness of Boletaceae (Wu *et al*., 2016; Sato & Toju, 2019). Although the extrinsic factors which contributed to this Miocene gene innovation remain to be investigated, the substantial differences in the intrinsic biology and host/habitat preferences may be relevant to the different histories of Suillineae, Sclerodermatineae and Boletineae.

In Agaricales, Russulales, Thelephorales and Pezizales the transition from saprotrophy to the symbiotic ecology coincided with the loss of most hydrolytic enzymes acting on lignocellulose (Kohler *et al*., 2015; Hess *et al*., 2018; Murat *et al*. 2018; Miyauchi *et al*., 2020; Looney *et al*., 2021; Marqués-Gálvez *et al*., 2021). Our analyses of the largest set of ectomycorrhizal Boletales genomes to date support the view that transitions from brown-rot to symbiosis entailed the widespread losses of PCWDEs acting on lignin, cellulose, hemicellulose and pectins. We found striking genomic signatures related to (hemi)cellulose- and lignin degradation genes that separate ectomycorrhizal species from their brown-rot relatives. Although brown-rot species are depauperate in PCWDEs compared to white-rots, their PCWDE repertoire, such as secreted GHs and polysaccharide lyases, is still substantially higher than the phylogenetically-related symbiotrophic species. Compared to their saprotrophic cousins, all ectomycorrhizal Boletales lack the set of enzymes required for efficient cellulose degradation consisting of cellobiohydrolases from families GH6 and GH7, and cellulose-binding module CBM1 appended to cellulases and endoglucanases. The absence of the synergistically acting GH7 and GH6 enzymes, acting on cellulose reducing and non-reducing ends, is a hallmark of ectomycorrhizal species in Agaricales and Russulales (Kohler *et al*., 2015; Wolfe *et al*. 2012; Looney *et al*., 2021). Concerning hemicelluloses, neither the endo-β-1,4-xylanases (GH11) acting on the xyloglucan backbone nor the debranching enzymes working synergistically with GH11 were found. Within the general evolutionary trend of PCWDEs loss, there is, however, a more specific dynamics at play. Different ectomycorrhizal fungi retain distinct sets of PCWDEs, suggesting that over their evolutionary history, symbiotic Boletales have become functionally diverse. For example, *Ch. virens* has a set of PCWDEs and FCWDEs closer to its brown-rot cousins. This likely reflects their ecological niche and variable dependence to their host plants. The smaller PCWDE repertoire was found in ectomycorrhizal Sclerodermatineae, such as *Pi. tinctorius* and *Pi. microcarpus*, reinforcing their dependence on the plant host. This may be related to their preferred ecological niche (i.e., sandy soils) with a scarce content in organic matter.

Despite their ecology, genomes of most symbiotic Boletales encode polygalacturonases (GH28, GH43), an endoglucanase GH5_5 with a CBM1 module, endoglucanases GH12, and one or two copies of xyloglucan-specific endo-β-1,4-glucanases (GH45) in *B. edulis*, *X. badius*, *Phlebopus* and *Suillus* species. The ß-1,4 endoglucanase (GH5_5-CBM1) is the ortholog of the *Laccaria bicolor* LbGH5-CBM1 involved in cell wall remodeling during the formation of the Hartig net and is an important determinant for successful symbiotic colonization of the *Laccaria bicolor*– *Populus tremula* × *alba* association (Zhang *et al*., 2018). The remaining set of GHs may thus play a role in host root colonization. In addition, the gene copy number of several CAZyme families involved in LOX (laccases, alcohol oxidases, glyoxal oxidases, benzoquinone reductases, LPMOs (AA9, AA14)) is similar in brown-rot and ectomycorrhizal species, suggesting that some symbiotic Boletales are capable of mild lignocellulose decomposition (e.g., litter bleaching) to scavenge nitrogen trapped in SOM as suggested by Floudas *et al*. (2020).

Ectomycorrhizal fungi are not all depauperate in PCWDEs. A few species, such as *Acephala macrosclerotium* (Leotiomycetes), have kept a substantial set of PCWDEs which are repressed in ectomycorrhizal root tips (Miyauchi *et al*., 2020). They may represent transitional steps from pure saprotrophy toward pure ectomycorrhizal symbiosis (Lebreton *et al*., 2021). Within the Boletales studied here, *Phlebopus* species in the Boletinellaceae appear to have a dual lifestyle. These boletes are able to establish ectomycorrhizal root tips with *Pinus kesiya* (Sanmee *et al*., 2010; Kumla *et al*., 2016), their basidiocarp ^13^C/^15^N isotopic signature is typical of ectomycorrhizal species (Cao *et al*., 2015) and we showed in the present study that their PCWDE repertoire is amongst the lower of the symbiotic boletes, supporting their adaptation to the symbiotic lifestyle. On the other hand, they are known to produce basidiocarps in African and Australian grasslands (Ji *et al*., 2011). The formation of fruiting bodies from soil free-living mycelium away from any known ectomycorrhizal host plants is in favor of a substantial saprotrophic ability of the mycelium to sustain the development of the sexual organs. Here, we confirmed that an ectomycorrhiza-forming *P. portentosus* mycelium can produce fruiting bodies on a mixed sawdust/seed substrate and we proposed that the activity of their limited set of PCWDEs and FCWDEs, such as laccases and chitinases, is sufficient to release the carbohydrates required for basidiocarp development in the absence of glucose from the plant. This mild decay mechanism may play a role in litter decomposition in natural settings in the absence of host trees.

Novel and recently-evolved genes, including effector-like MiSSPs, are thought to be responsible for the specific attributes of individual mycorrhizal lineages (Kohler *et al*., 2015; Martin *et al*., 2016). However, we have shown by using a phylostratigraphic analysis that a large proportion of symbiosis-related genes are orthologous to genes encoded by saprotrophic lineages that arose long before the evolution of the mutualistic associations (Miyauchi *et al*., 2020). These genes encoded by saprotrophic ancestors have been co-opted for the symbiotic lifestyle. They are related to key ecological traits, such as N and P acquisition (e.g., organic N- and P-degrading secreted enzymes, nutrient transporters) already present in free-living saprotrophic ancestors of symbiotrophic species. Unfortunately, the identification of symbiosis-related genes in boletes, such as *B. edulis*, is precluded by our inability to produce ectomycorrhizas *in vitro* for boletes; these late-stage symbionts being poor colonizers of tree seedlings. We therefore used symbiosis-induced genes identified in transcript profiling from *Pa. involutus, Pi. tinctorius*, *Pi. microcarpus* and *S. luteus* (Kohler *et al*., 2015) and assessed their sequence conservation/divergence within the Boletales genomes. We confirmed that most of the symbiosis-induced genes are conserved amongst saprotrophic and symbiotic Boletales, and only a small set (∼10%) of symbiosis-induced transcripts (encoding MiSSPs or proteins of unknown function) are species-specific. This proportion is much higher (> 30 %) in the Agaricales (Miyauchi *et al.,* 2020).

In conclusion, the genomes of symbiotrophic Boletales species have a greatly increased content in TEs and a much reduced set of genes coding for PCWDEs by comparison with their brown-rot relatives. Besides, several species in the Boletaceae, Paxillaceae and Boletinellaceae have kept a substantial set of endoglucanases and LPMOs acting on cellulose/hemicellulose and fungal polysaccharides indicating that they may partly decompose SOM by a combined activity of oxidative and hydrolytic enzymes as shown for *P. involutus* and *Laccaria bicolor* (Nicolás *et al*., 2019). In the latter ectomycorrhizal symbionts, the host tree actively controls the decomposing activities of the associated fungi by controlling the amount of photosynthetic carbon provided to the fungus. These genomic features are shared with ectomycorrhizal Agaricales and Russulales (Miyauchi *et al*., 2020; Looney *et al*., 2021). It is tempting to speculate that TEs accelerated the evolutionary rate of genes encoding effector-like small secreted proteins, proteases, and lipases. Here, we also showed that ectomycorrhizal Boletaceae, to a smaller extent Suillaceae, experienced an obvious expansion of gene families in the late Miocene, an evolutionary event possibly contributing to the evolutionary success of boletes. This study produced the most inclusive phylogenomic and genomic analyses for the Boletales order to date, including characterization of several transitions from brown-rot to ectomycorrhizal lifestyles. Although further studies on a larger sample of Boletales genomes are required to substantiate our results and conclusions, the present study provides novel insights on our understanding of the mechanisms that influence the evolutionary diversification of boletes, the symbiosis evolution and the molecular interactions with their host plants.

## ACKNOWLEDGEMENTS

This work was supported by the Strategic Priority Research Program of Chinese Academy of Sciences (grant number XDB31000000) (to WG), the Laboratory of Excellence ARBRE (grant number ANR-11-LABX-0002-01), the Beijing Advanced Innovation Center for Tree Breeding by Molecular Design, the Region Lorraine and the European Regional Development Fund (to F.M.M.). We also acknowledge grants from the International Partnership Program of Chinese Academy of Sciences (grant number 151853KYSB20170026 to Z.L.Y.), the National Natural Science Foundation of China (grant number 31970015 to G.W.), the Yunnan Ten Thousand Talents Program Plan Young & Elite Talents project (to G.W.), the Youth Innovation Promotion Association of the Chinese Academy of Sciences (grant number 2017436 to G.W.), the Yunling Scholars Funds of the Ten-Thousand-Talents Program of Yunnan Provincial Government (to Z.L.Y.), the Momentum Program of the Hungarian Academy of Sciences (grant number LP2019/13-2019 to L.G.N.), the National Research, Development and Innovation office (contract number GINOP-2.3.2-15-2016-00052 to L.G.N.) and the Natural Science Foundation of Yunnan Province (grant number 2017FB025 to Y.C.). This work was also funded by the U.S. Department of Energy Joint Genome Institute, a DOE Office of Science User Facility, and supported by the Office of Science of the U.S. Department of Energy under Contract No. DE-AC02-05CH11231 within the framework of the Mycorrhizal Genomics Initiative (CSP # 305), Metatranscriptomics of Forest Soil Ecosystems project (CSP # 570) and the 1000 Fungal Genome project (CSP # 1974). GW would like to thank the China Scholarship Council for supporting his research stay at INRAE and thank Dr. Hong Luo for his advices to perform genomic DNA isolation.

## AUTHOR CONTRIBUTIONS

F.M.M. conceived and coordinates the Mycorrhizal Genomics Initiative. F.M.M., G.W. and Z.L.Y. designed the project. G.W., S.M. and F.M.M. wrote the manuscript with the help of L.G.N. and T.V.. I.V.G. coordinated genome sequencing and annotation at JGI. G.W., B.F., Z.L.Y., Y.C. and J.X. coordinated genome and transcriptome sequencing, and annotation at KIB. A. Kuo, C.D., J.J., H.H., K.L., V.N. performed transcriptome sequencing, assembly and gene annotation at JGI. E.D. and B.H. performed CAZyme annotations. S.M., G.W., E.M. and F.M.M. performed comparative genome analyses. G.W., S.M. and A. Kohler carried out the RNA-seq analysis. T.V. and L.G.N. reconstructed and analyzed genome-wide duplication and contraction events. I.G.C., H.P., J.M.P., J.M. and J.W.S. provided genomes used in this study.

## AVAILABILITY OF DATA AND MATERIALS

Genome assemblies and gene annotations for JGI-sequenced Boletales used in this study are available via the JGI fungal genome portal MycoCosm (see the Boletales portal at https://mycocosm.jgi.doe.gov/boletales/boletales.info.html). Sequenced genomes are also available at the National Center for Biotechnology Information (NCBI) GenBank (see Table S1 for accession codes/BioProjects). The complete transcriptome data sets are available at the Gene Expression Omnibus at NCBI (http://www.ncbi.nlm.nih.gov/geo/). All other data supporting the findings of this study are included within the article and its additional files.

## CODE AVAILABILITY

Visual omics ShingoTools - PRINGO, TINGO, SynGO, VINGO are available at GitHub: https://github.com/ShingoMiyauchi

The program COMPARE is at GitHub: https://github.com/laszlognagy/COMPARE.

## Supporting Information

**Fig. S1.** The coverage (a) and copy number (b) of transposable elements (TEs) identified in Boletales and selected genomes.

**Fig. S2.** Bar plot showing the scaffold size of selected fungi in their phylogenetic order.

**Fig. S3.** The percentage of syntenic blocks in selected Boletales genomes.

**Fig. S4.** Genome macrosynteny in Boletineae and allied Boletinellaceae.

**Fig. S5.** Locations of CAZyme-coding genes in scaffold 1 of *Boletus edulis* (Boled5) with corresponding syntenic regions in other allied Boletales fungi.

**Fig. S6.** Locations of CAZyme-coding genes on scaffold 2 of *Boletus edulis* (Boled5) with corresponding syntenic regions in other allied Boletales fungi.

**Fig. S7.** Locations of CAZyme-coding genes on scaffold 5 of *Boletus edulis* (Boled5) with corresponding syntenic regions in other allied Boletales fungi.

**Fig. S8.** Locations of SSP genes on scaffold 2 of *Boletus edulis* (Boledp1) with corresponding syntenic regions in other allied Boletales fungi.

**Fig. S9.** Presence of CAZyme-coding genes in scaffolds of *Boletus edulis* (Boled5) aligned with other Boletales fungi.

**Fig. S10.** Ordination analysis assessing the correlation between TE coverage and selected protein-coding gene counts.

**Fig. S11.** Conservation of the protein-coding genes framing the CBM1-GH6 cellobiohydrolase gene located on scaffold 2 of the saprotrophic *Hydnomerulius pinastri* (Hydpi2) and missing from the symbiotrophic Boletales species.

**Fig. S12.** Gene conservation and innovation in Boletales fungi.

**Fig. S13.** Diversification of the protein-coding gene repertoires in Boletales families.

**Fig. S14.** Number of secreted and total CAZyme-coding genes in the genomes.

**Fig. S15.** Non-metric multidimensional scaling (NMDS) analysis of total (right panel) and secreted (left panel) CAZyme-coding gene repertoires in Boletales and other species.

**Fig. S16.** Number of genes coding for proteases and protease inhibitors (according to the MEROPS database) in the genomes of 28 Boletales and selected outgroup species.

**Fig. S17.** The transcription level of *Ph. portentosus* PP33 genes for plant and fungal cell wall degrading enzymes under three conditions.

**Fig. S18.** The transcription level of *Ph. portentosus* PP33 CAZyme-coding genes under three conditions.

**Fig. S19.** Phylogenetic conservation of symbiosis-induced genes in Boletales.

**Fig. S20.** Number of genes coding for secondary metabolism, transcription factors and membrane transporters.

**Table S1.** Strain information of the newly sequenced Boletales species.

**Table S2.** Taxonomic affiliation, genomic features and statistics for the 34 included genomes.

**Table S3.** Data for PERMANOVA on genome features of Boletales species.

**Table S4.** Pair-wise PERMANOVA and PERMANOVA of genome features.

**Table S5.** The statistics of generalized least squares analysis with the Brownian motion model.

**Table S6.** Identified transposable elements in the Boletales species and outgroupss.

**Table S7.** CAZy count in top 10 scaffolds and total scaffolds of the 11 species with high-quality genomes out of 15 target fungi.

**Table S8.** Mean distances between repeat elements and genes.

**Table S9.** PERMANOVA results among TEs and SSPs, GT2, GT.

**Table S10.** Core, dispensable, specific genes of 34 fungi.

**Table S11.** Count of EcM-specific and brown-rot-specific genes in 28 Boletales species.

**Table S12.** COMPARE function enrichment.

**Table S13.** Data for Kirisame (drizzle) plot illustrating SSPs in syntenic regions with SSP annotations.

**Table S14.** *Phlebopus portentosus* PP33 transcription levels for genes coding for PCWDEs+FCWDEs and all CAZymes.

**Supporting information-Methods.**

