## Supplementary material for "Evolutionary innovations through gain and loss of genes in the ectomycorrhizal Boletales": Supporting information-Figures-6Sept2021.docx

**Supporting information: Supplemental figures**

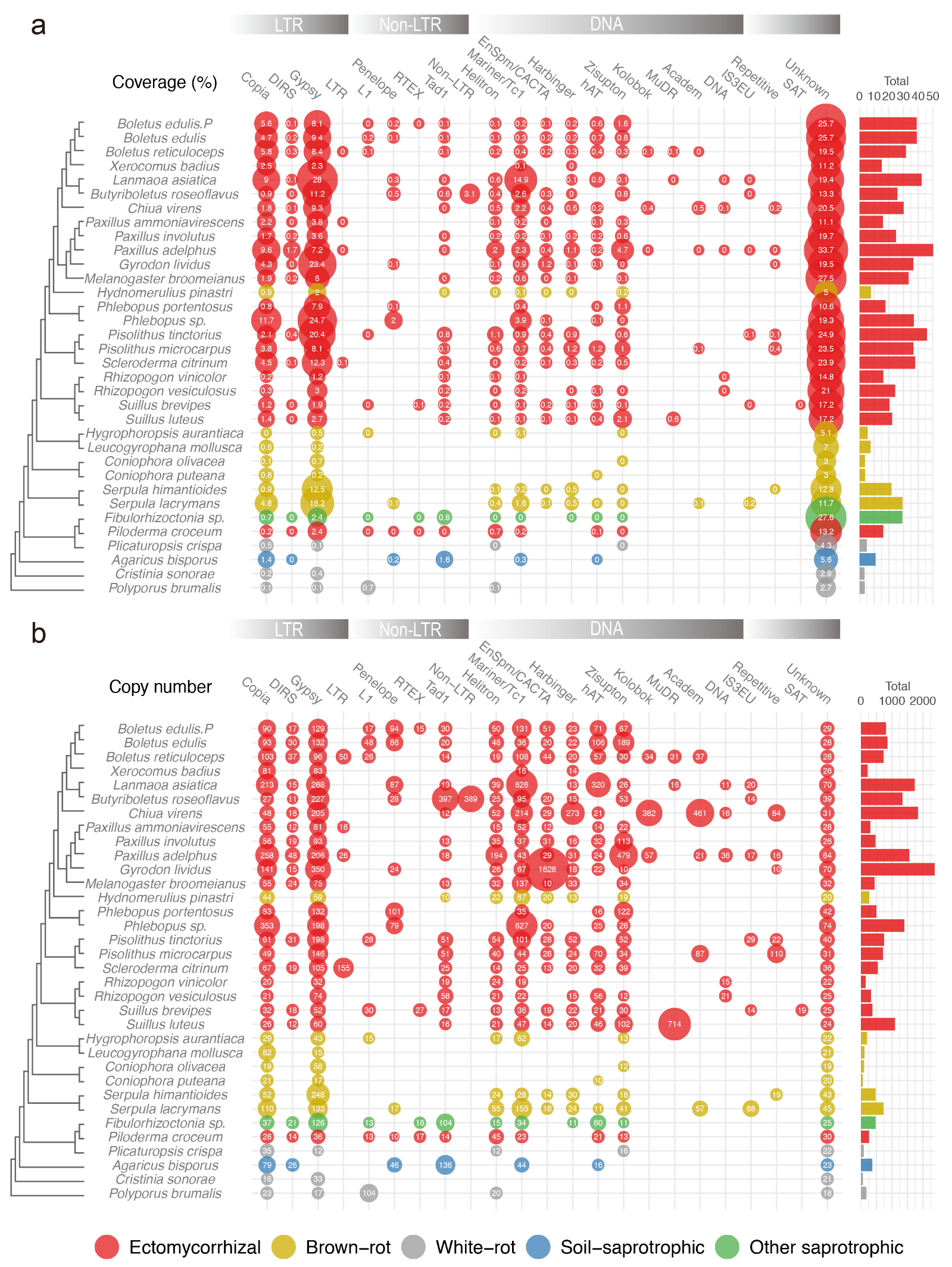

**Fig. S1.** **The coverage (a) and copy number (b) of transposable elements (TEs) identified in Boletales and selected genomes.** LTR, long terminal repeat retrotransposons; non-LTR, non-long terminal repeat retrotransposons; DNA: DNA transposons; repetitive/SAT: simple repeats; unknown: unclassified repeated sequences. The bubble size is proportional to the coverage (a) / copy number (b) of each of TE (showed inside the bubbles). The bars show the total coverage per genome. See also Support Information-Table S2 and Table S6.

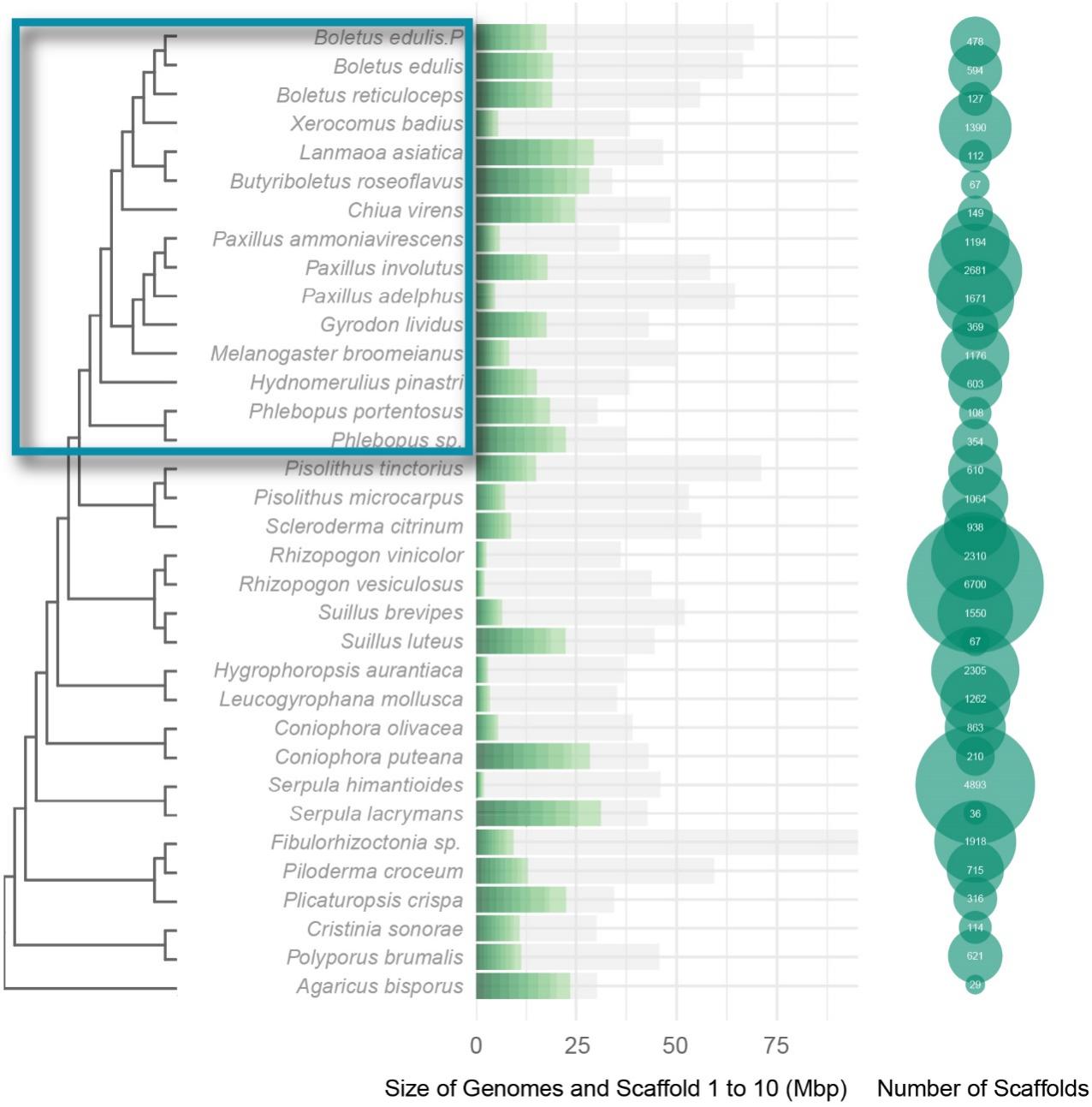

**Fig. S2.** **Bar plot showing the scaffold size of selected fungi in their phylogenetic order**. Green bars, top ten largest scaffolds. Grey bars, total scaffold size. A total of 11 (out of 15) fungi with good genome assemblies were selected for the synteny analysis (green frame). *X. badius* (Xarba1), *P. ammoniavirescens* (Paxam1), *P. adelphus* (Paxru1) and *M. broomeianus* (Melbro1) were not included in the analysis due to their highly fragmented genomes.

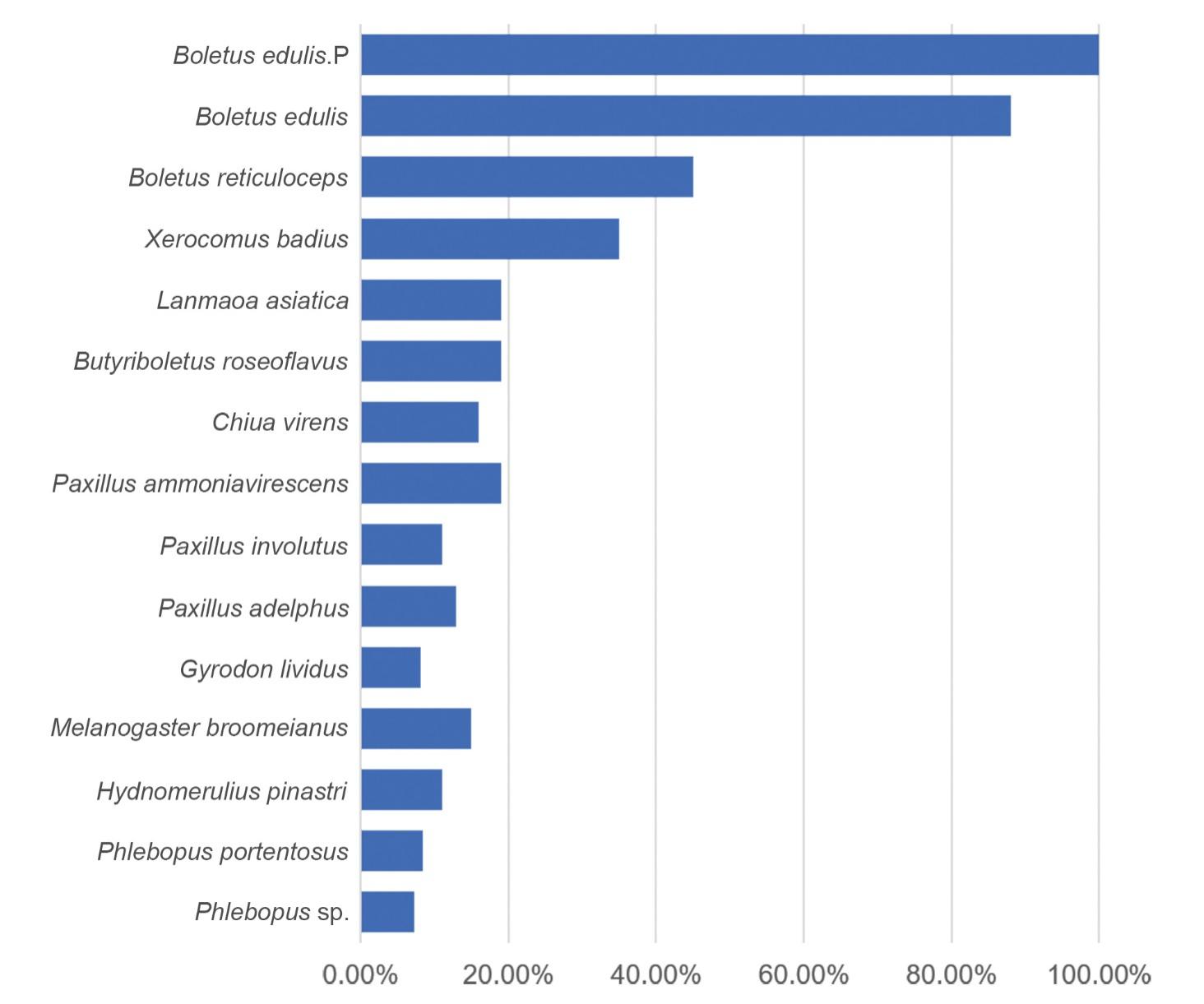

**Fig. S3.** **The percentage of syntenic blocks in selected Boletales genomes.** Synteny was determined using *Boletus edulis* (Boled5) as the reference species.

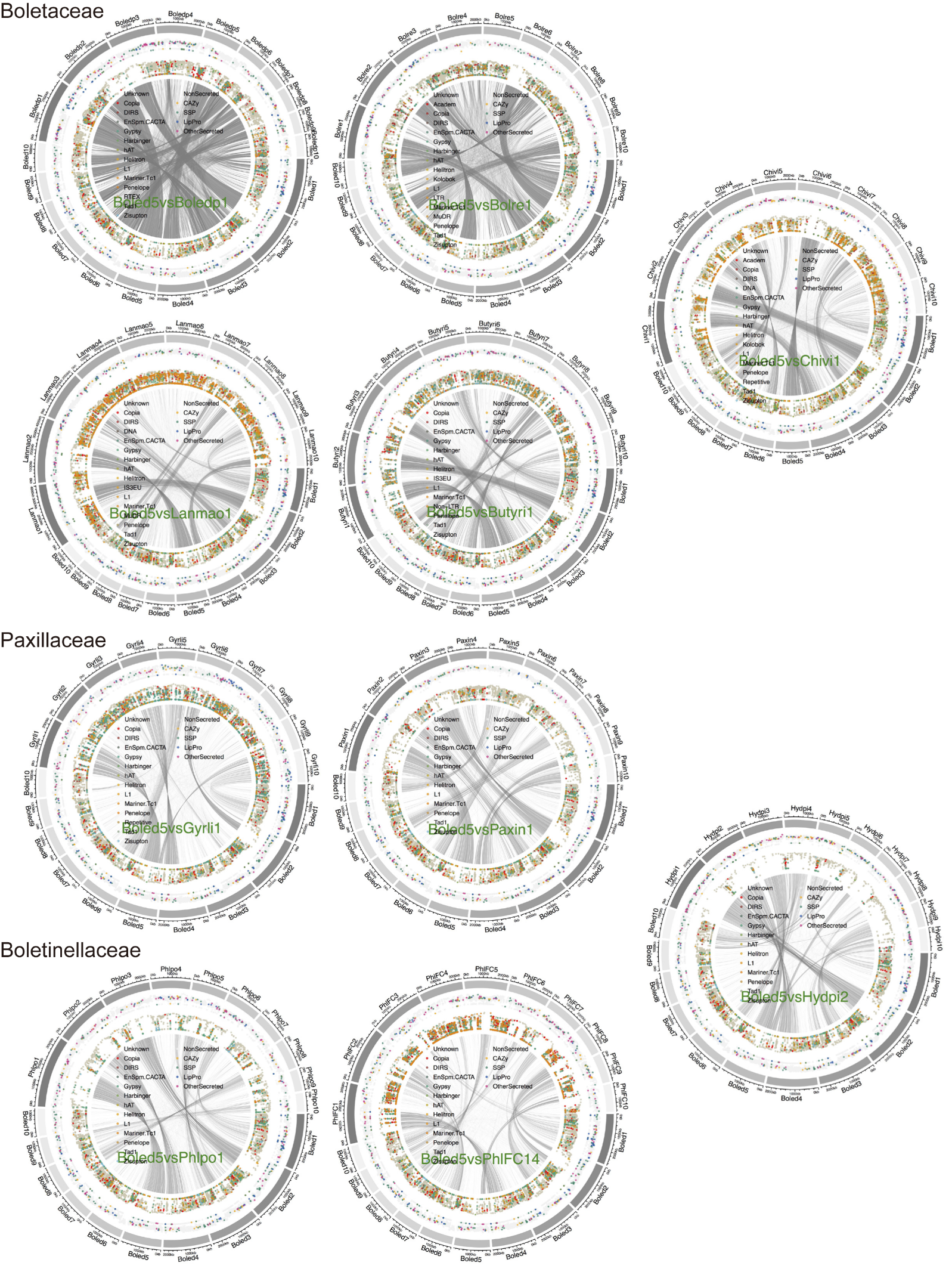

**Fig. S4.** **Genome macrosynteny in Boletineae and allied Boletinellaceae.** Hanabi (firework) plots show genes for intracellular and extracellular proteins with TEs. Synteny was identified using *Boletus edulis* (Boled5) as the reference genome. Grey links indicate syntenic regions between two species. Outer circle, scaffold size; first inner circle, genes coding for secreted CAZymes, SSPs, lipases and proteases (see legends inside circles). Second inner circle, TEs (see legends inside circles). Vertical axis of inner circles, mean distances of neighbouring genes/repeats; short distances between the genes/repeats result in dots towards the centre of plots, whereas long distances result in dots towards the outer circle.

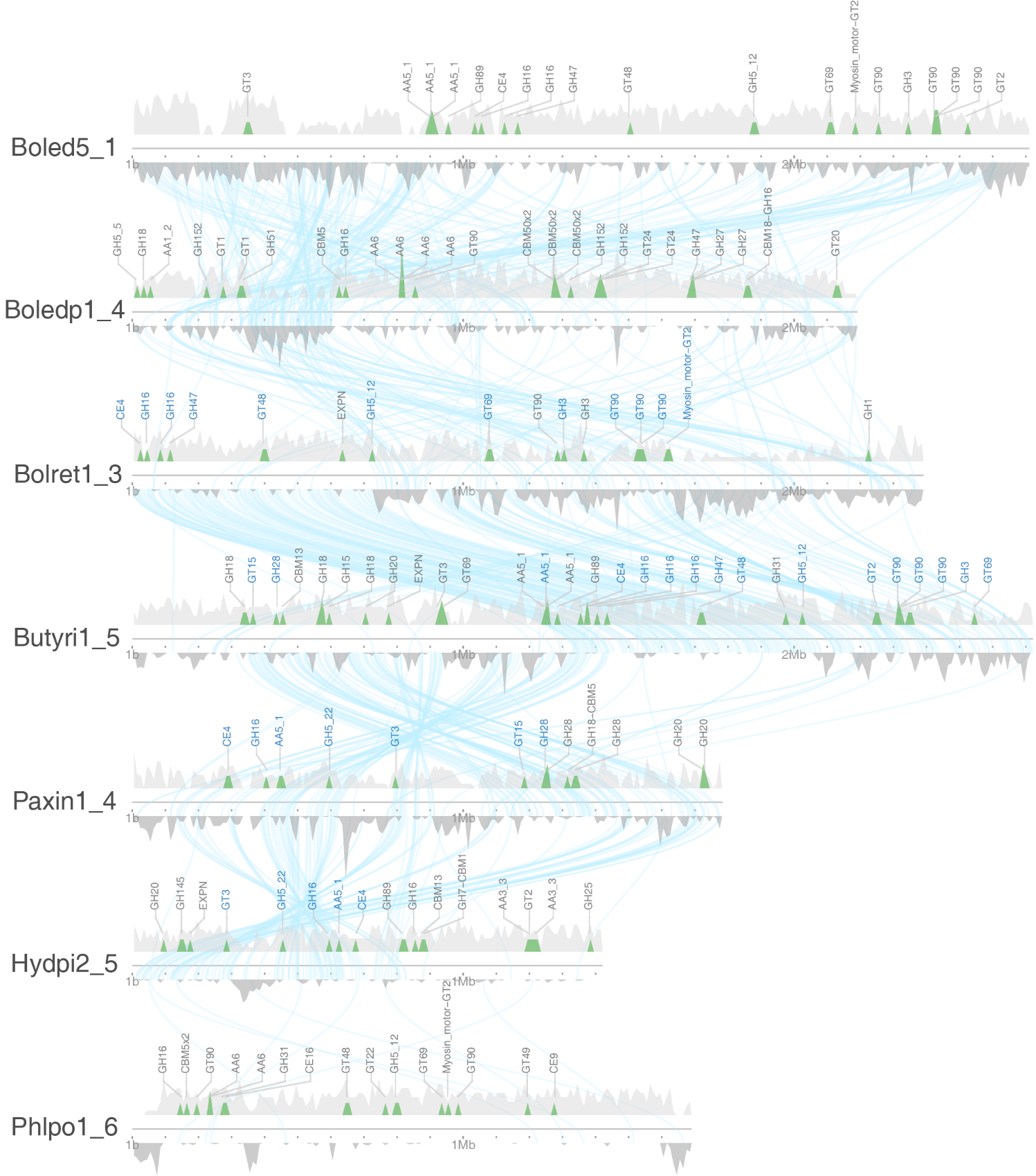

**Fig. S5. Locations of CAZyme-coding genes in scaffold 1 of *Boletus edulis* (Boled5) with corresponding syntenic regions in other allied Boletales fungi.** Kirisame (drizzle) plot illustrates CAZyme coding genes and repeat elements in syntenic regions. Blue lines: Syntenic regions. Red: Genes in common among the species compared. Blue: Genes shared with neighbouring species. Orange: Newly introduced genes in synteny evolved from the common ancestor. Upward peaks (light grey): Density of all genes. Downward peaks (Dark grey): Density of repeat elements including transposable elements and unknown repeats. Species are in evolutionary order. Shortened JGI fungal identification with scaffold number are shown on the left side of the scaffolds. (see Support Information-Table S2 for corresponding species names). See Support Information-Fig. S9 for a summary of CAZymes.

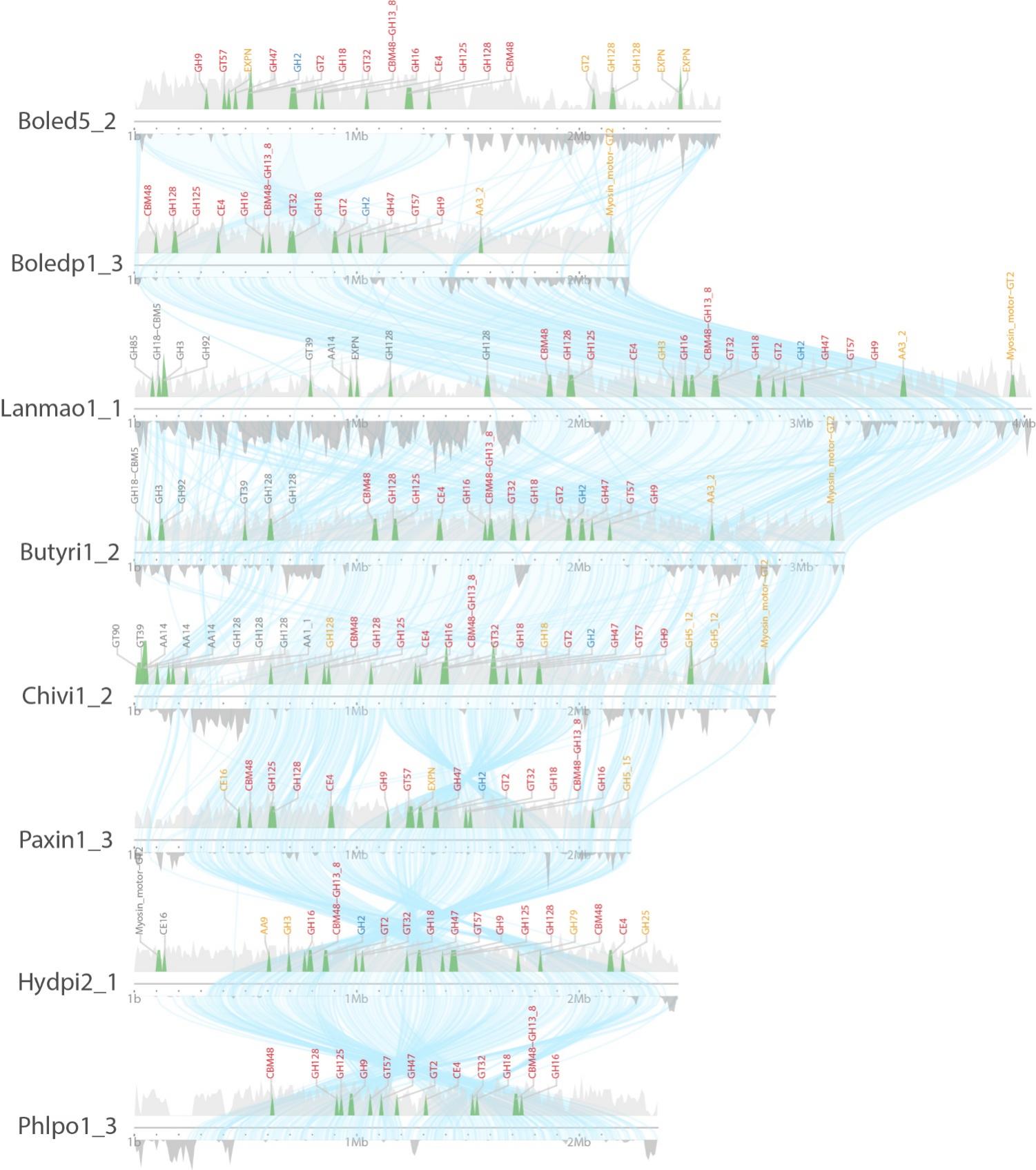

**Fig. S6.** **Locations of CAZyme-coding genes on scaffold 2 of *Boletus edulis* (Boled5) with corresponding syntenic regions in other allied Boletales fungi.** Kirisame (drizzle) plot shows the location of CAZyme genes and repeated elements in syntenic regions. Blue lines, Syntenic regions; Red fonts, CAZyme genes in common among the species; Blue fonts, CAZyme genes shared with neighbouring species; Orange font, newly inserted CAZyme genes in the scaffold and evolving from the common ancestor. Upward peaks (light grey): Density of protein-coding genes. Downward peaks (Dark grey): Density of repeat elements including TEs and unknown repeats. Species are stacked according to their phylogenetic relationship. JGI assembly identification with scaffold number are shown on the left side of the scaffolds (see Support Information-Table S2 for corresponding species names). See Support Information-Fig. S9 for a summary of CAZymes.

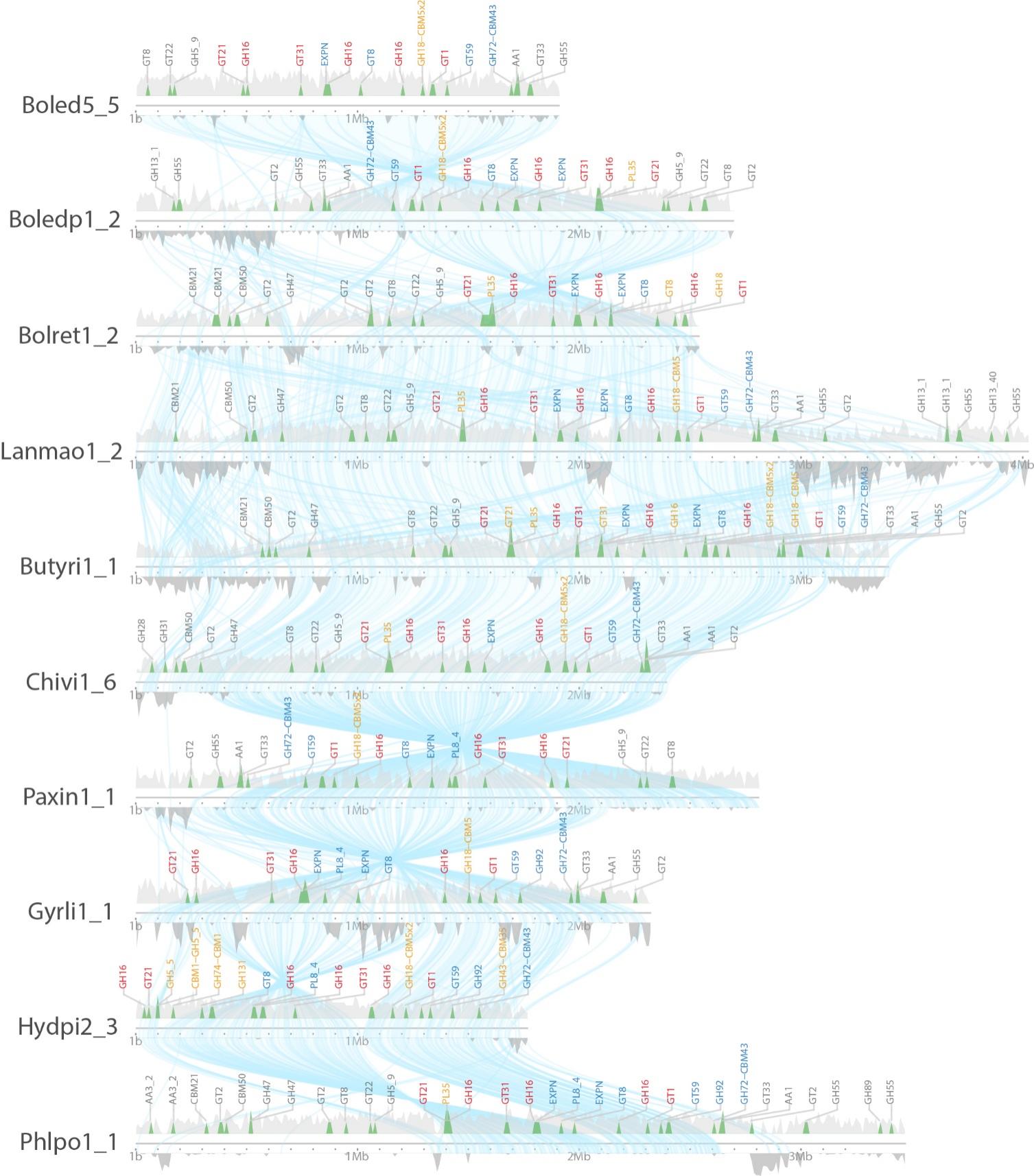

**Fig. S7. Locations of CAZyme-coding genes on scaffold 5 of *Boletus edulis* (Boled5) with corresponding syntenic regions in other allied Boletales fungi.** Kirisame (drizzle) plot shows the location of CAZyme genes and repeated elements in syntenic regions. Blue lines, Syntenic regions; Red fonts, CAZyme genes in common among the species; Blue fonts, CAZyme genes shared with neighbouring species; Orange font, newly inserted CAZyme genes in the scaffold and evolving from the common ancestor. Upward peaks (light grey): Density of protein-coding genes. Downward peaks (Dark grey): Density of repeat elements including TEs and unknown repeats. Species are stacked according to their phylogenetic relationship. JGI assembly identification with scaffold number are shown on the left side of the scaffolds (see Support Information-Table S2 for corresponding species names). See Support Information-Fig. S9 for a summary of CAZymes.

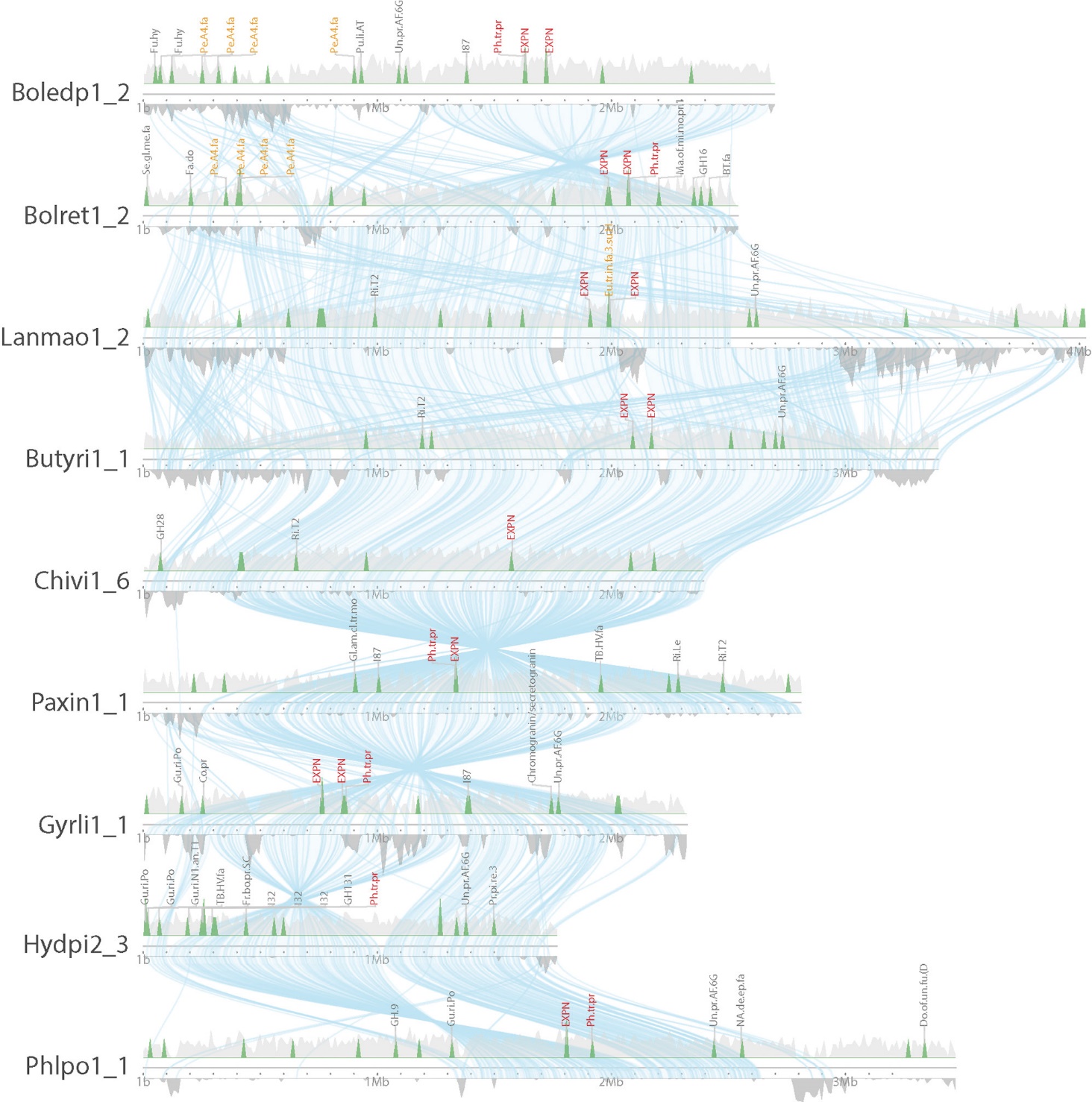

**Fig. S8. Locations of SSP genes on scaffold 2 of *Boletus edulis* (Boledp1) with corresponding syntenic regions in other allied Boletales fungi.** Kirisame (drizzle) plot show the location of SSP genes and repeat elements in syntenic regions. Gene names in red, genes in common; Gene names in orange, newly inserted genes in the syntenic region. Upward peaks (light grey), density of all protein-coding genes; downward peaks (dark grey), density of repeat elements including TEs and unknown repeats. Blue links, syntenic regions. Species are stacked according to their phylogenetic relationship. JGI assembly identification with scaffold number are shown on the left side of the scaffolds (see Fig. 1 for corresponding species names). Abbreviations: Pe.A4.fa: G01; Aspergillopepsin-2, Peptidase_A4_family; Eu.tr.in.fa.3.su.H, Eukaryotic translation initiation factor 3 subunit H; JAB1/Mov34/MPN/PAD-1, ubiquitin protease; EXPN, Expansin; Ph.tr.pr, Phosphatidylglycerol/phosphatidylinositol transfer protein. See Support Information-Table S13 for details.

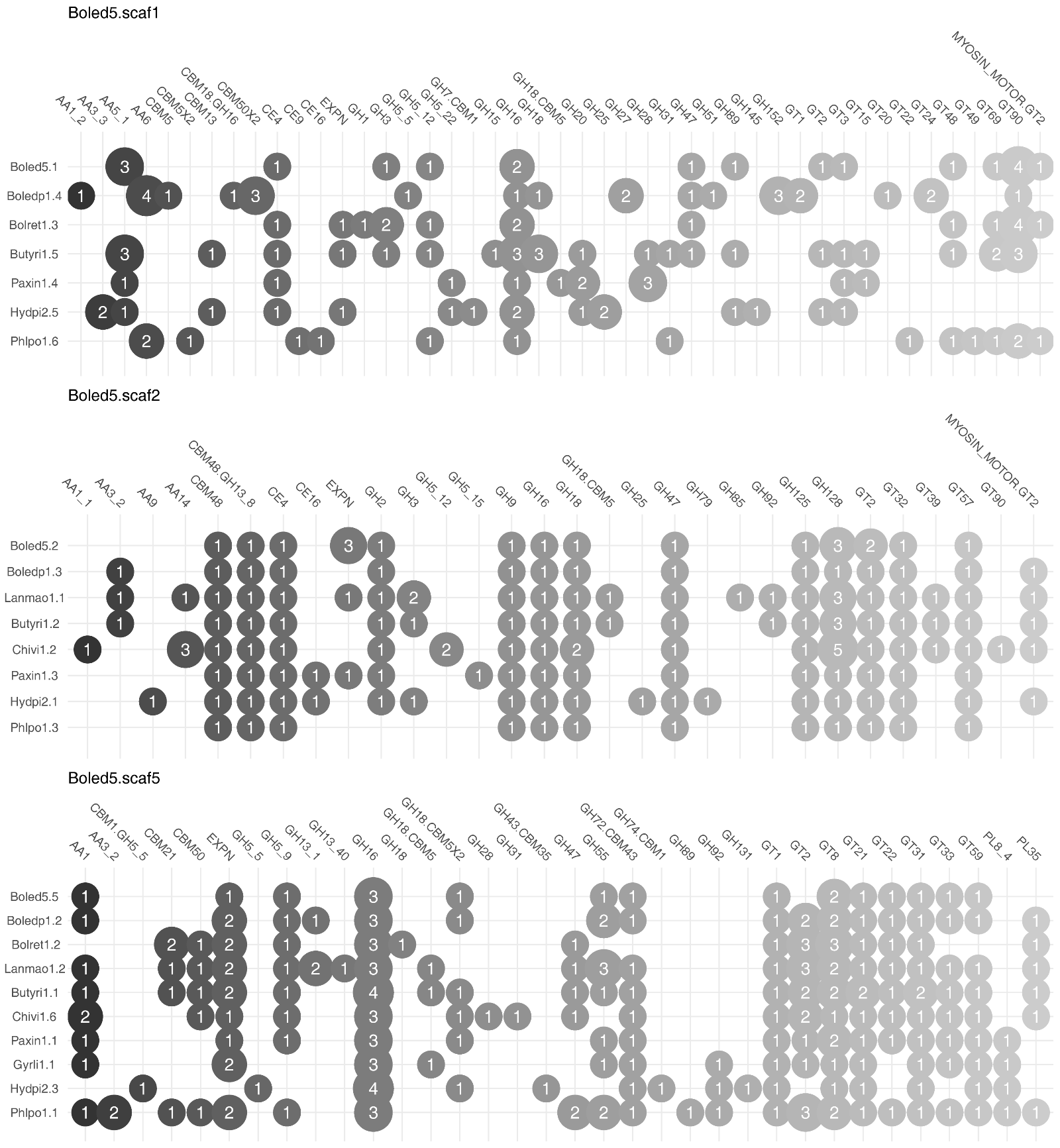

**Fig. S9. Presence of CAZyme-coding genes in scaffolds of *Boletus edulis* (Boled5) aligned with other Boletales fungi.** A bubble plot shows scaffold 1, 2, and 5 of *B. edulis* with corresponding syntenic regions of other allied Boletales fungi. Bubble size corresponds to the gene copy number.

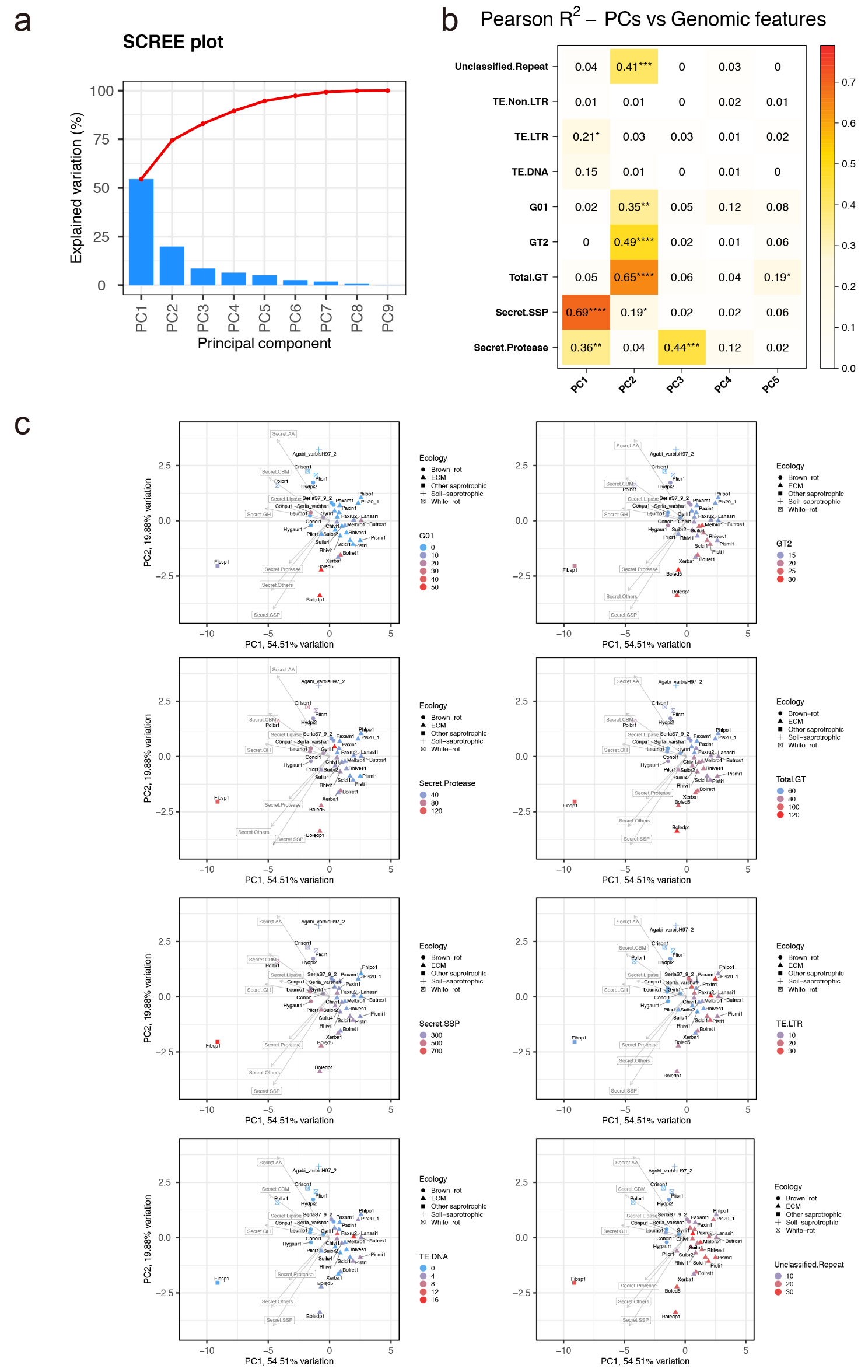

**Fig. S10. Ordination analysis assessing the correlation between TE coverage and selected protein-coding gene counts.** a) Scree plot showing the level of principal component variance captured from the gene count data. PC 1 to 3 explained over 80% of total PCA variance. b) Correlations of principal components with the selected genomic features. The values are Pearson R squared. Significant p values are indicated (FDR adjusted p < 0.0001:****, 0.001:***, 0.01:**,0.05:*). c) Overlaid genomic features on two dimensional coordinates of 34 fungi based on PC1 and PC2. Shapes represent ecological groups of fungi (white-rot fungi, brown-rot fungi, soil/litter saprotrophs, and ectomycorrhizal fungi). Arrows show five highest loadings of variables driving variation in PCs. The genomic features include the genomic coverage of DNA transposons, LTR- and non-LTR retrotransposons, unclassified repeats, the gene counts for protease G01, chitin synthase GT2, total glycosyl transferases (gT), SSPs (Secret.SSPs) and secreted proteases (Secret.Protease).

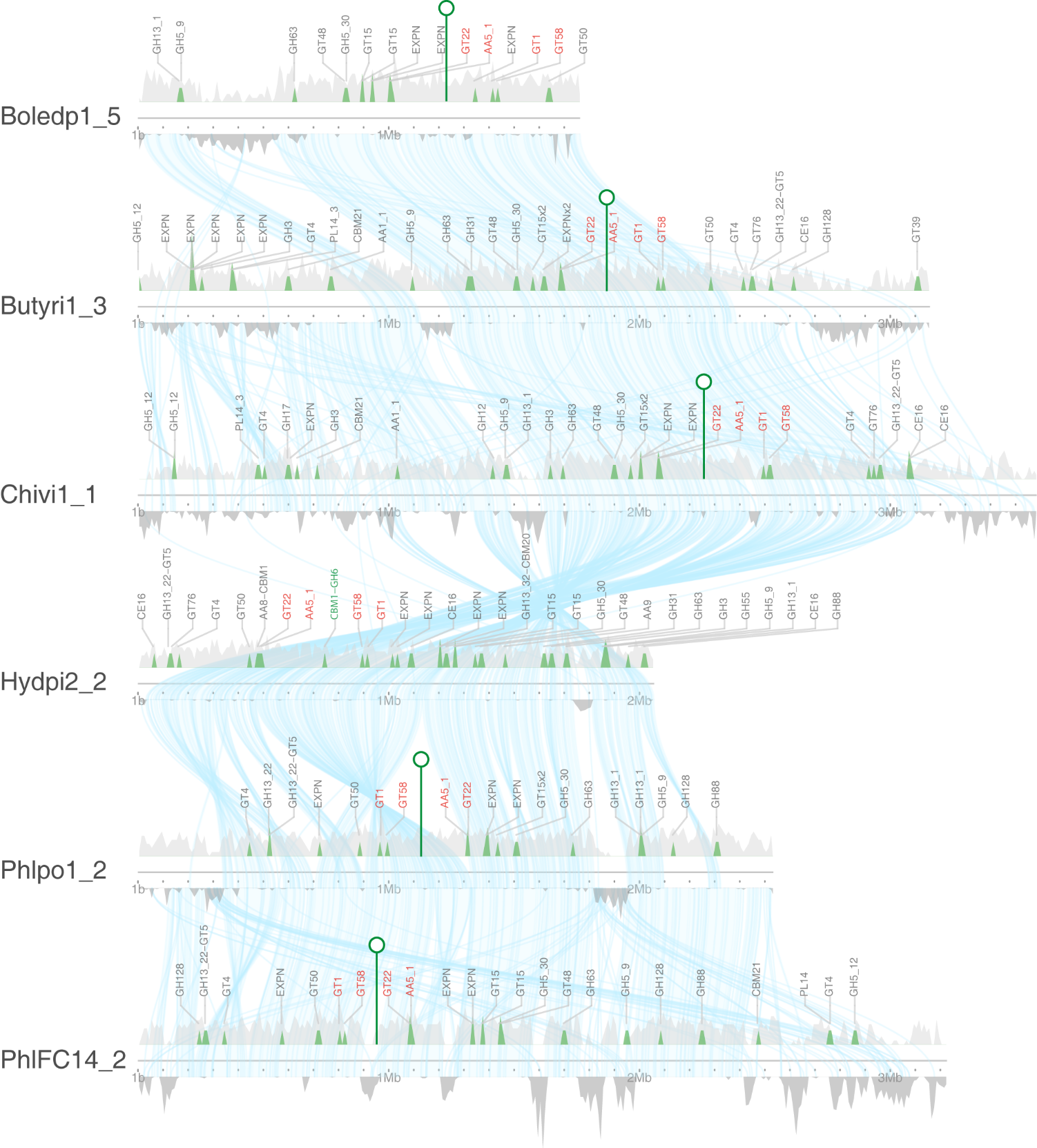

**Fig. S11.** **Conservation of the protein-coding genes framing the *CBM1-GH6* cellobiohydrolase gene located on scaffold 2 of the saprotrophic *Hydnomerulius pinastri* (Hydpi2) and missing from the symbiotrophic Boletales species.** The Kirisame (drizzle) plot shows CAZyme genes and repeat elements in syntenic regions. The location of the *CBM1-GH6* gene in *H. pinastri* is marked by a green triangle and green font. The corresponding location for the missing *CBM1-GH6* gene in symbiotic boletes is indicated by a pin-like tag. Conserved CAZyme genes framing the *CBM1-GH6* are in red. Upward peaks (light grey), density of all genes; downward peaks, dark grey, density of repeat elements including TEs and unknown repeats. Blue links, syntenic regions. Species are stacked according to their phylogenetic relationship. JGI assembly identification with scaffold number are shown on the left side of the scaffolds (see Support Information-Table S2 for corresponding species names).

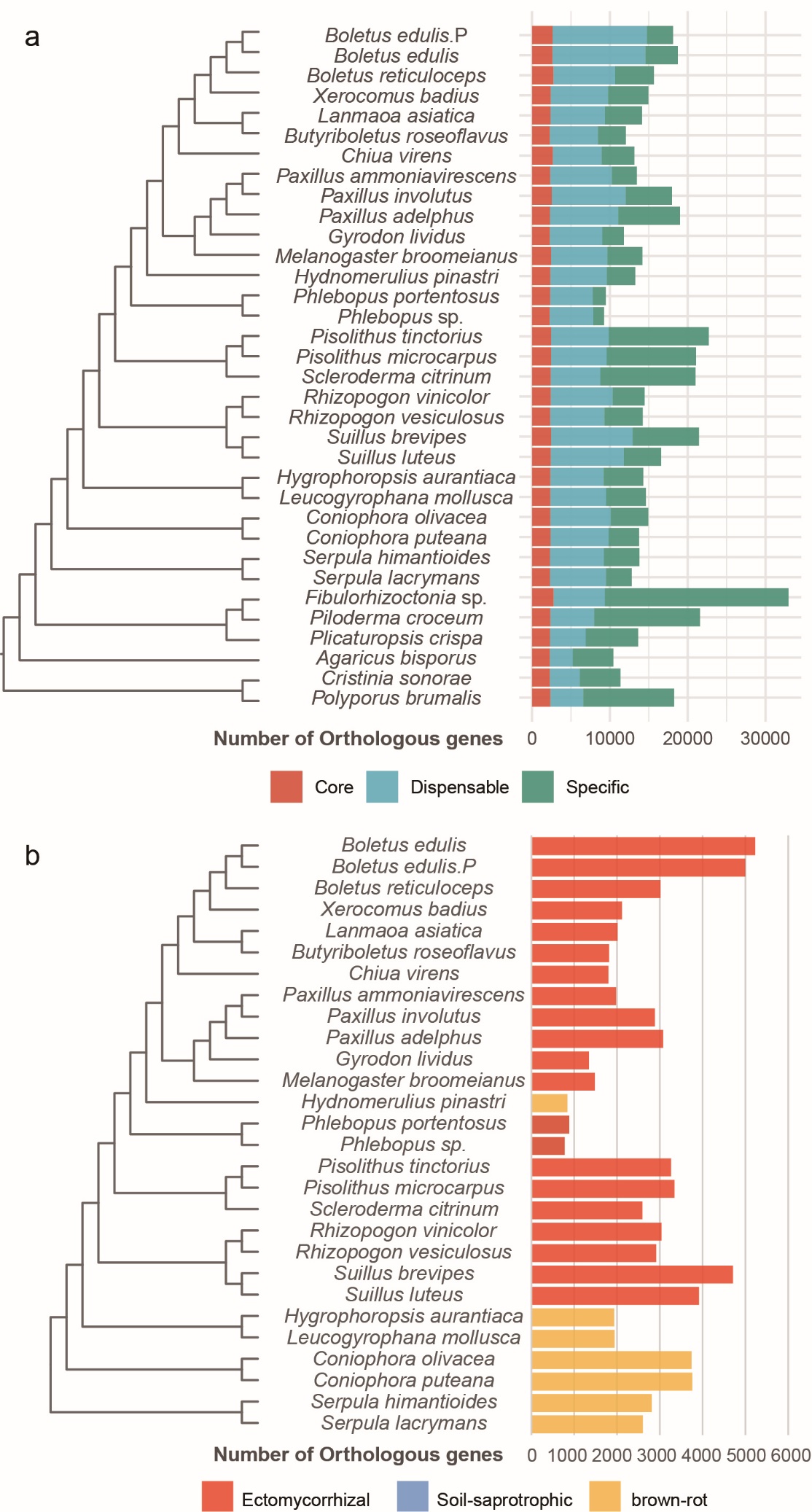

**Fig. S12.** **Gene conservation and innovation in Boletales fungi.** a) Stack bars stand for conserved proteins shared among species (in red), dispensable proteins (in blue), and species-specific (in green); b) sets of conserved proteins shared among ectomycorrhizal Boletales (in red) or shared among brown-rot Boletales (in yellow). The number of orthogroups for each category are listed in Support Information-Table S10 and Table S11.

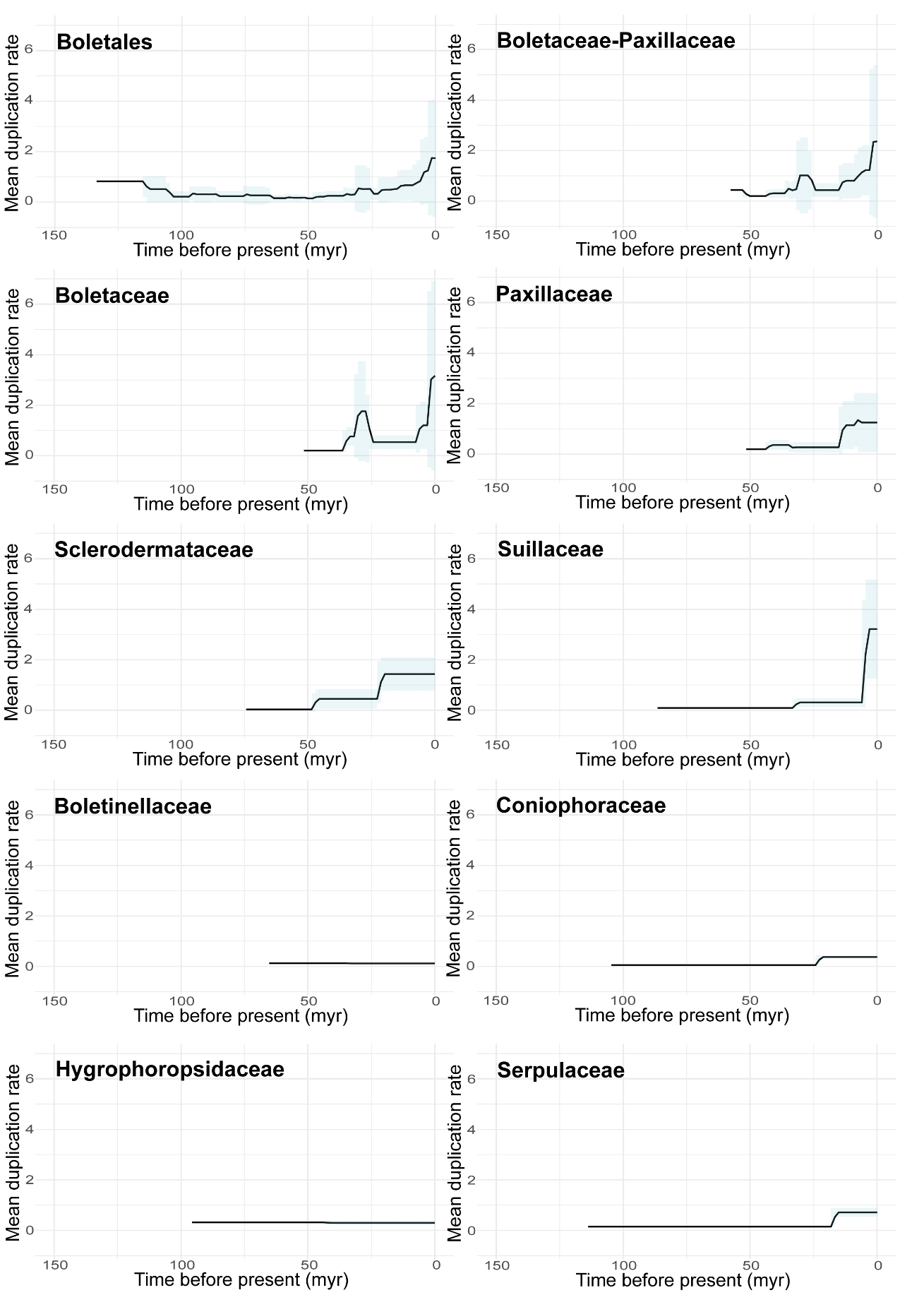

**Fig. S13.** **Diversification of the protein-coding gene repertoires in Boletales families.** Plots depict the gene duplication rates through time reflecting the evolution of gene repertoires in different clades of Boletales. Solid black line represents the mean of the gene duplication rate across lineages and time. Blue bars display the standard deviation of the gene duplication rates in a given time frame. Time frames were calculated by dividing the time interval of Boletales evolution over 150 million years (Myr) by 100.

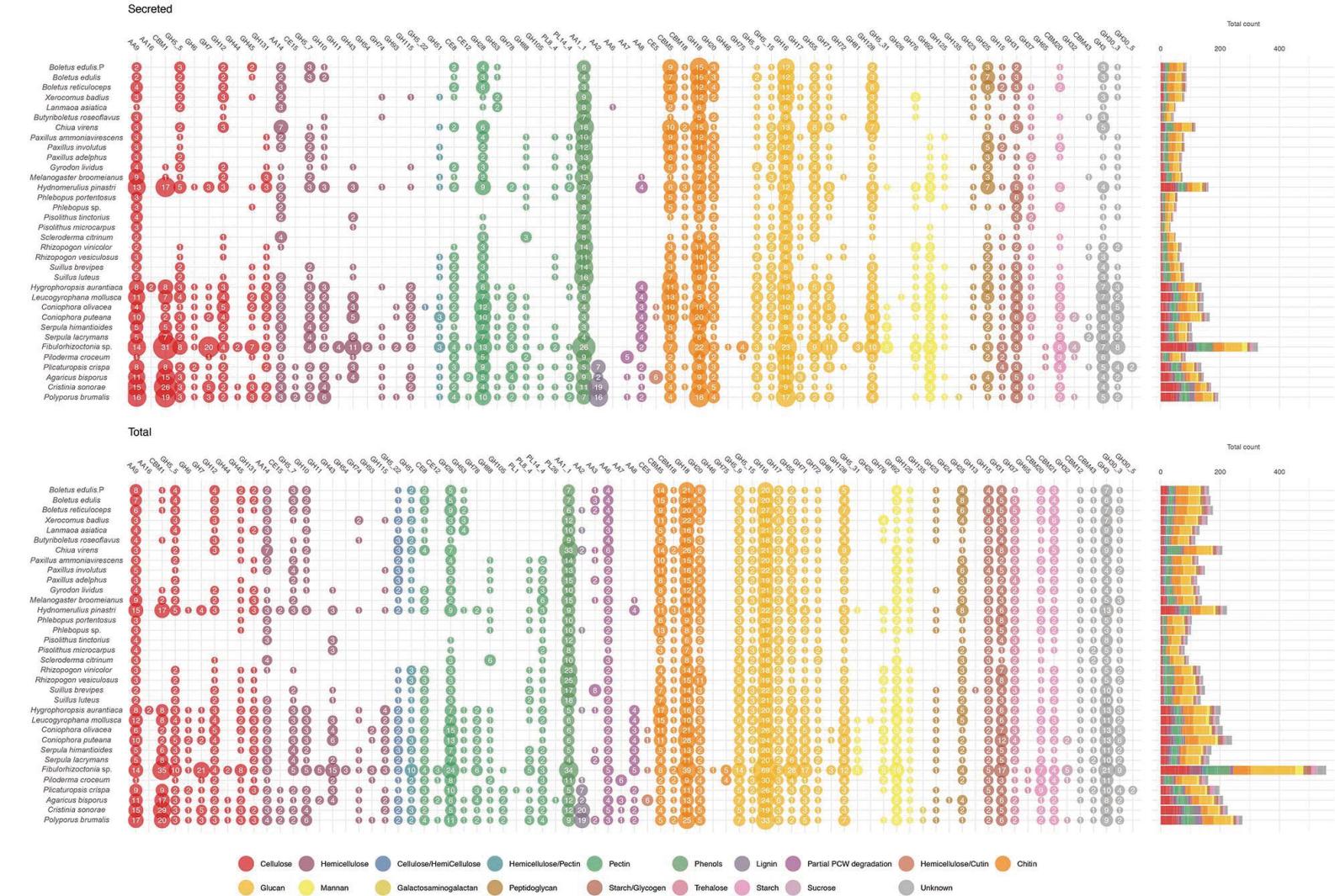

**Fig. S14.** **Number of secreted and total CAZyme-coding genes in the genomes.** All the CAZyme domains are counted (i.e. secreted and non-secreted). The fungal species are color-coded according to their ecology. The potential substrate, i.e. cellulose, hemicellulose, lignin, and pectin (plant cell walls); chitin, glucan, mannan (fungal cell walls) and peptidoglycan (bacterial cell walls), are color-coded, see at the bottom of the figure.

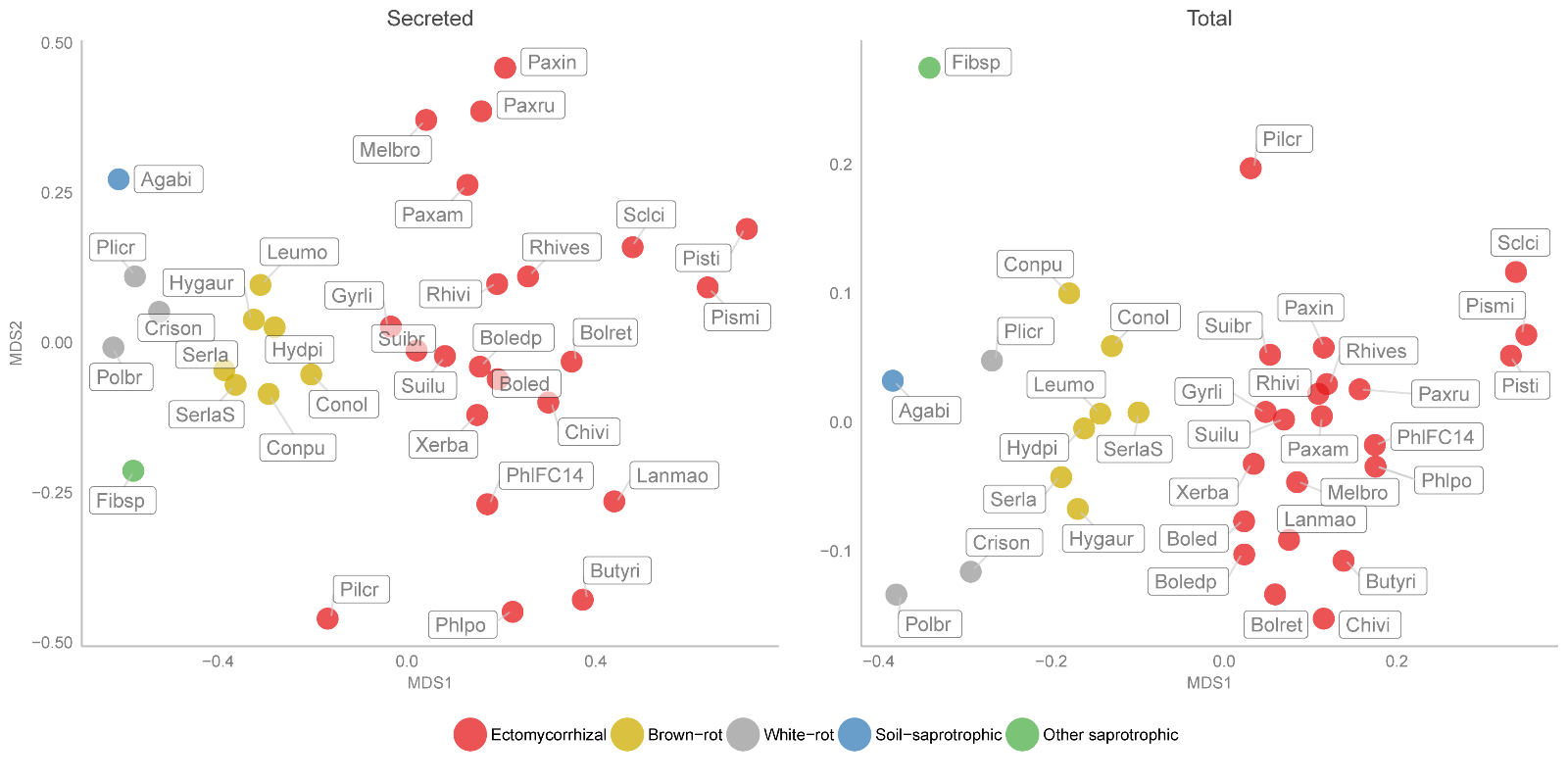

**Fig. S15.** **Non-metric multidimensional scaling (****NMDS) analysis of total (right panel) and secreted (left panel) CAZyme-coding gene repertoires in Boletales and other allied species.** The fungal species are color-coded based on their ecology and identified using the JGI assembly IDs.

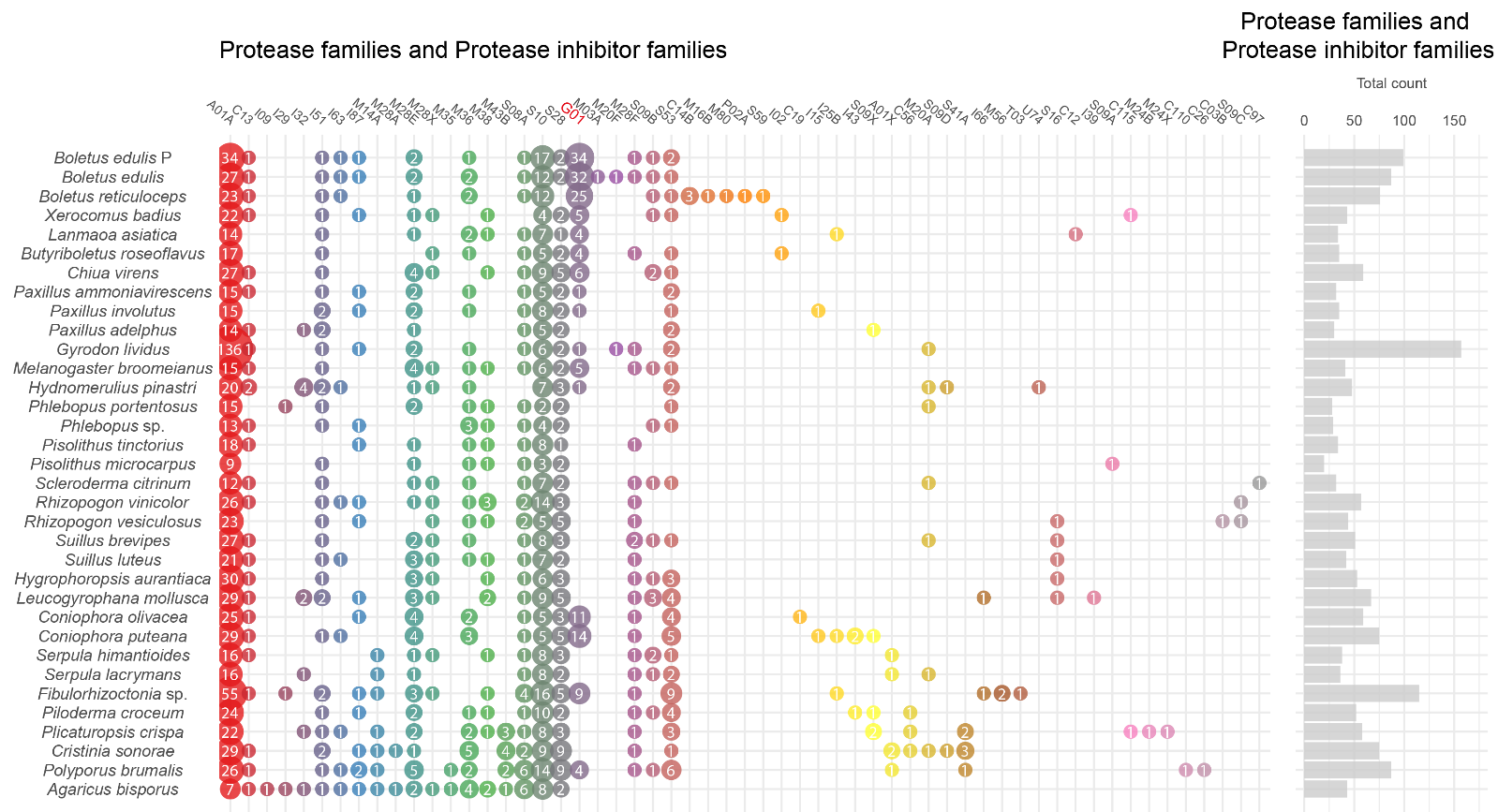

**Fig. S16.** **Number of genes coding for proteases and protease inhibitors (according to the MEROPS database) in the genomes of 28 Boletales and selected outgroup species.** Note a much higher gene copy number for the secreted pepstatin-insensitive carboxyl proteinase G01 (in purple) in *Boletus* species.

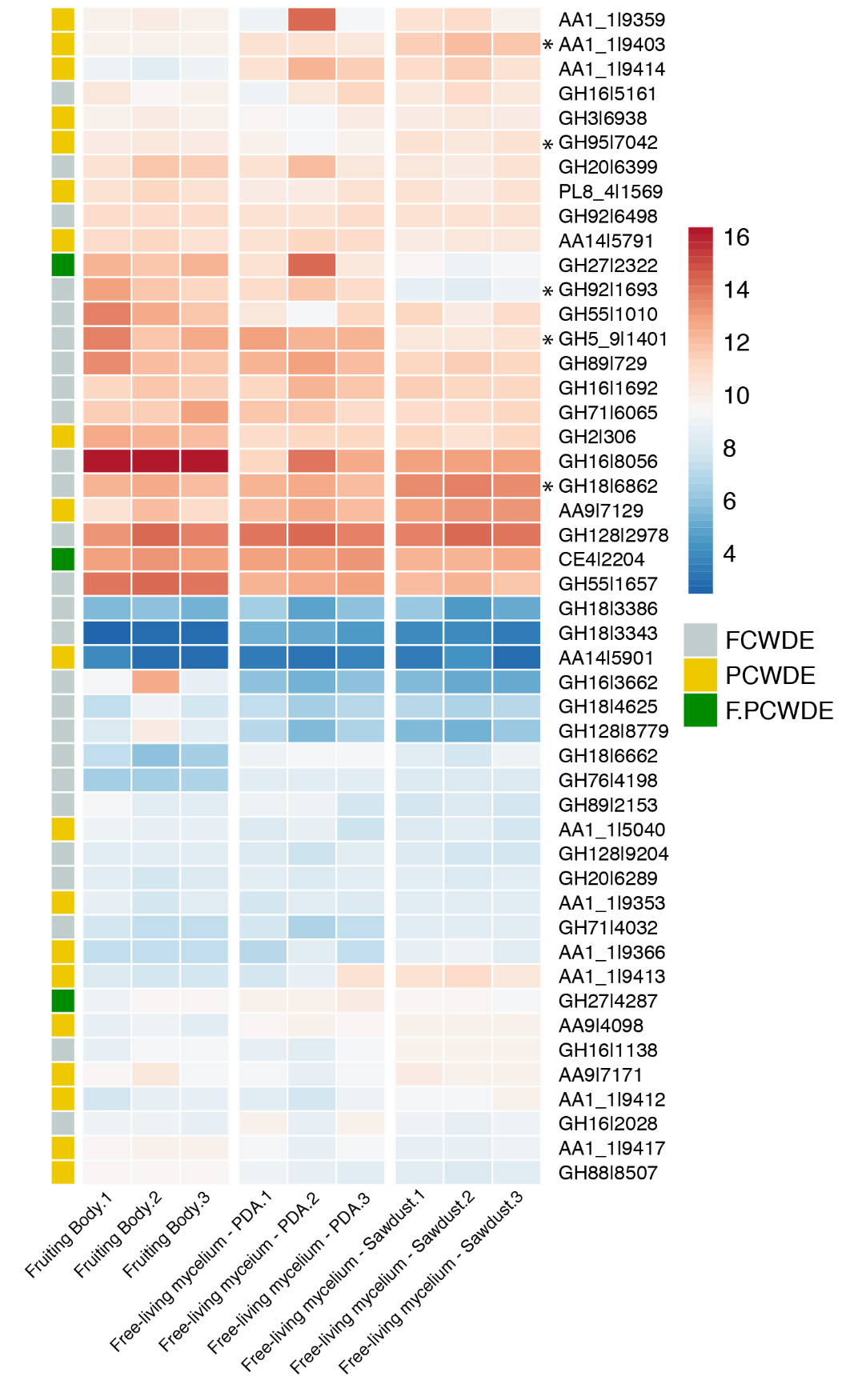

**Fig. S17.** **The transcription level of *Ph. portentosus* PP33 genes for plant and fungal cell wall degrading enzymes under three conditions.** Asterisks show significantly differentially expressed genes between the PDA and sawdust media (FDR adj p < 0.05). PCWDE: Plant cell wall degrading enzymes. FCWDE: Fungal cell wall degrading enzymes. F.PCWDE: CAZymes degrade plant and fungal cell walls. Gene expression in log2 is shown per sample. See Support Information-Table S14.

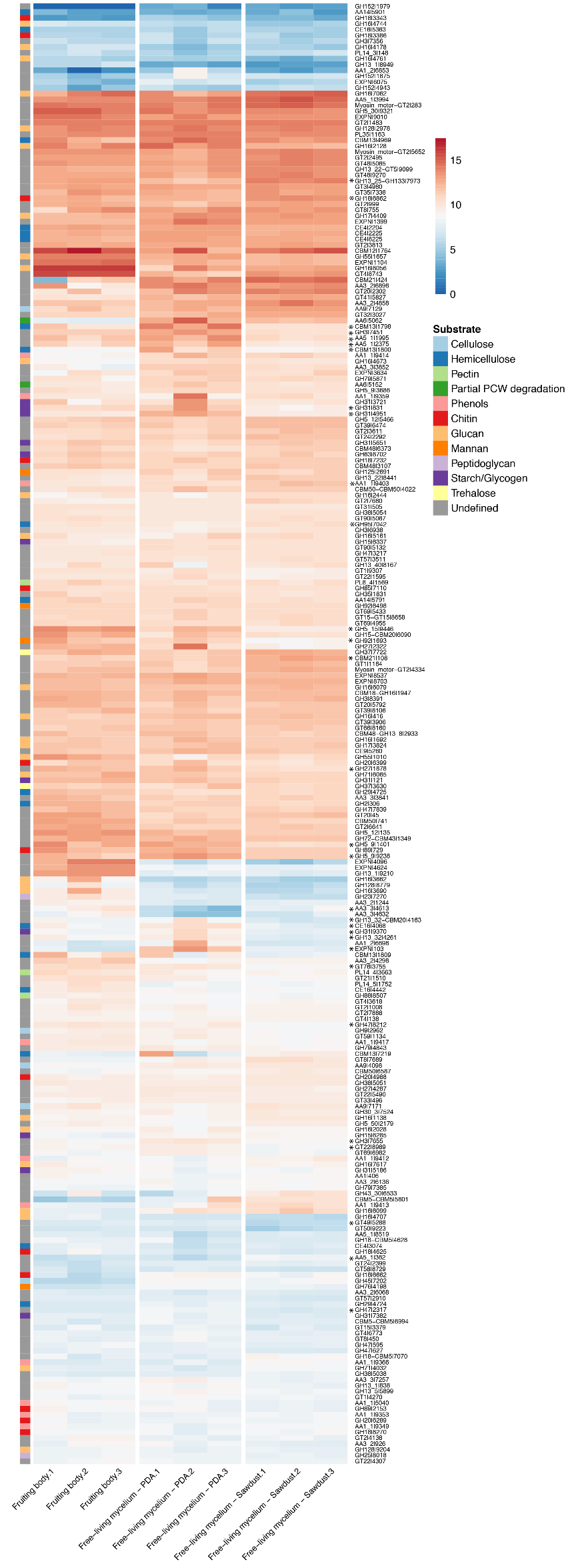

**Fig. S18.** **The transcription level of *Ph. portentosus* PP33 CAZyme-coding genes under three conditions.** Asterisks show significantly differentially expressed genes between the PDA and sawdust media (FDR adj p < 0.05). The vertical bar on the left shows the substrate specificity of CAZymes. Gene expression in log2 is shown per sample. See Support Information-Table S14.

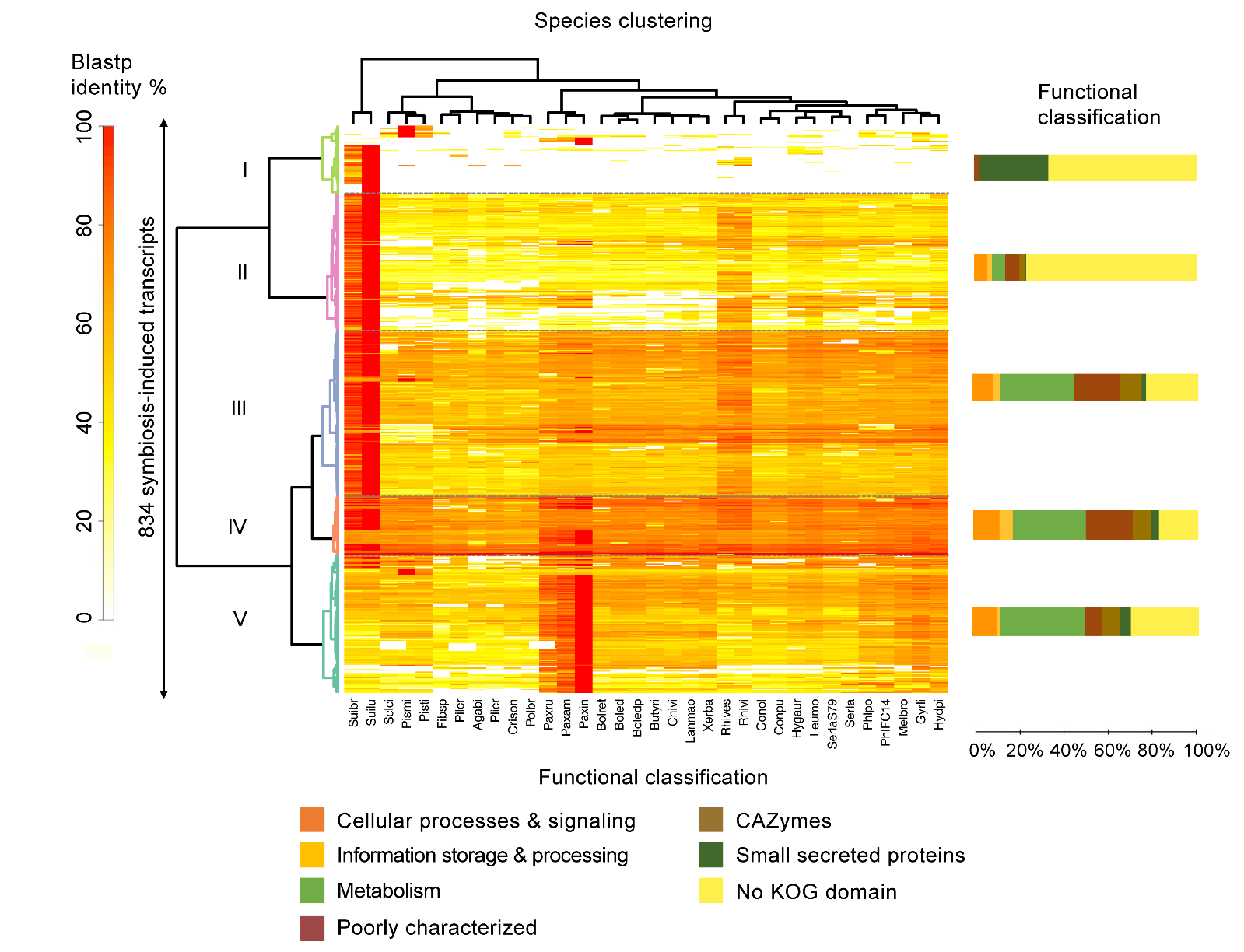

**Fig. S19.** **Phylogenetic conservation of symbiosis-induced genes in Boletales.** Heat maps display BLASTP sequence similarity of 275 ectomycorrhiza-induced proteins. The heat maps depict a double-hierarchical clustering of protein sequences encoded by symbiosis-upregulated genes (rows, fold change ≥ 5 in symbiotic tissues compared to free-living mycelium, false discovery rate-corrected P ≤ 0.05); right panel, functional categories (KOG) are given for each cluster of sequences in % as bargrams. Clusters I – VI correspond to group of sequences sharing the same level of protein sequence similarity based on BlastP. Data were visualized and clustered using R (package HeatPlus97). The hierarchical clustering was done by using a Euclidian distance and Ward clustering method. Color scale on the left (white to red) shows the % of sequence identity according to BLASTP. The symbiosis-induced genes were retrieved from the EcM transcriptomes of *Paxillus involutus, Pisolithus microcarpus*, *Pisolithus tinctorius* and *Suillus luteus*. For a description of the analysis, see Miyauchi et al. (2020).

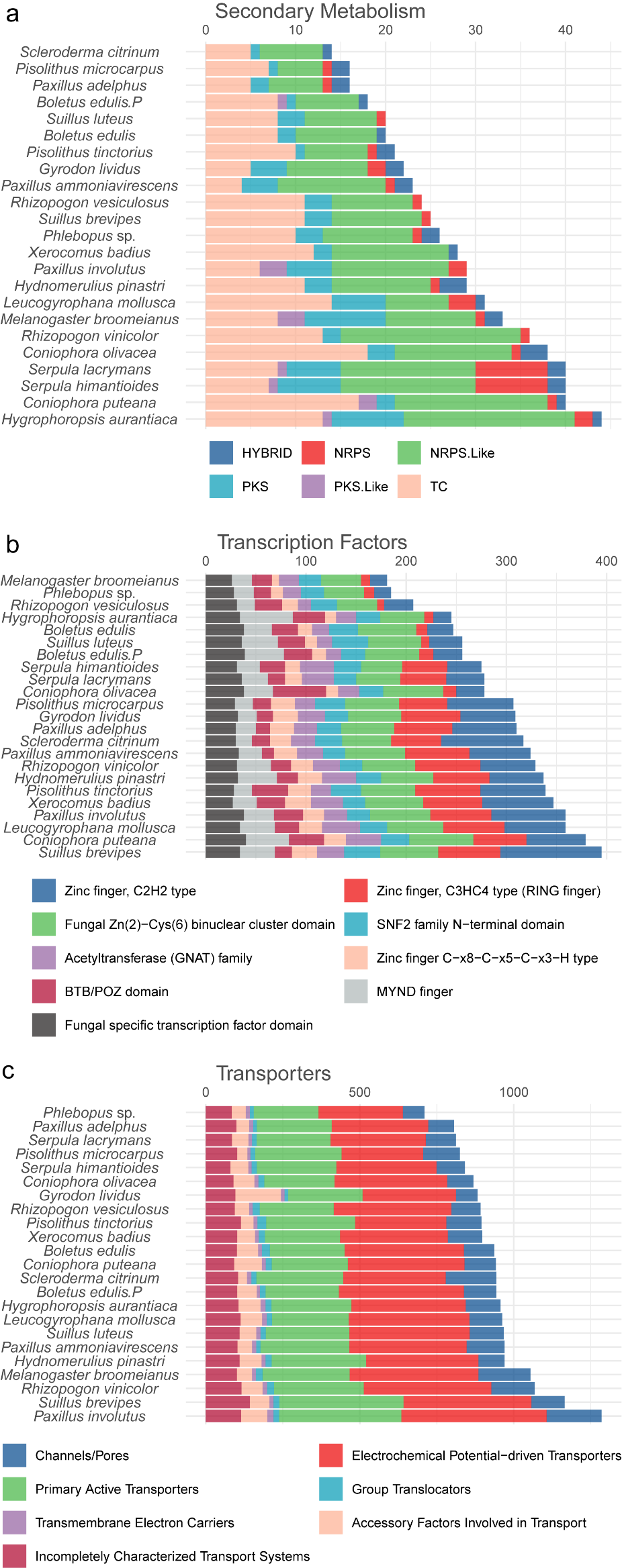

**Fig. S20.** **Number of genes coding for secondary metabolism, transcription factors and membrane transporters.** The gene annotation on JGI MycoCosm was used. (**a**) Biosynthetic gene clusters (BGC) coding for secondary metabolites. The count of BGC types per species are shown. HYBRID: PKS-NRPS mixed. NRPS: Non-ribosomal peptide synthetase cluster. NRPS.like: NRPS-like fragment. PKS: Polyketide synthase. PKS.like: Other types of PKS cluster TC: Terpene cyclases/synthases. DMAT: Dimethylallyl diphosphate synthase. (**b**) Transcription factors. (**c**) Membrane transporters.
