## Supplementary material for "Evolutionary innovations through gain and loss of genes in the ectomycorrhizal Boletales": Supporting information-Methods_rev-gwu-6Sept2021.docx

Gang Wu *et al.*

**Genome sequencing, assembly and annotation**

In this study, we compared the genomes of 28 fungal species of Boletales (*Boletus edulis* BED1*, Boletus edulis* Přilba*, Boletus reticuloceps, Butyriboletus roseoflavus, Chiua virens, Coniophora olivacea, Coniophora puteana, Gyrodon lividus, Hydnomerulius pinastri, Hygrophoropsis aurantiaca, Lanmaoa asiatica, Leucogyrophana mollusca, Melanogaster broomeianus, Paxillus adelphus, Paxillus ammoniavirescens, Paxillus involutus, Phlebopus portentosus, Phlebopus sp., Pisolithus microcarpus, Pisolithus tinctorius, Rhizopogon vesiculosus, Rhizopogon vinicolor, Scleroderma citrinum, Serpula himantioides, Serpula lacrymans, Suillus brevipes, Suillus luteus, Xerocomus badius)*, four species of the sister order Atheliales (*Cristinia sonorae*, *Fibulorhizoctonia psychrophila, Piloderma croceum (= P. olivaceum), Plicaturopsis crispa*) and two outgroup genomes (*Agaricus bisporus* (Agaricales), *Polyporus brumalis* (Polyporales)). Most of these genomes have been sequenced, assembled and annotated by using the Joint Genome Institute (JGI) pipelines methods and protocols. Below, we only provide the protocols used for the sequencing, assembly and gene prediction for the seven new Boletales genomes used in this comparative analysis.

The genome of *Hygrophoropsis aurantiaca, Leucogyrophana mollusca* and *Phlebopus* sp. FC_14 were sequenced at JGI. The genome of *Phlebopus* sp. FC_14 was sequenced using the Pacific Biosciences (PacBio) sequencing platform. One µg of genomic DNA was sheared to 10 kb using Covaris g-TUBE. The sheared DNA was treated with DNA damage repair mix followed by end repair and ligation of blunt adapters using SMRTbell Template Prep Kit 1.0 (Pacific Biosciences). The library was purified with AMPure PB beads. PacBio sequencing primer was then annealed to the SMRTbell template library and sequencing polymerase was bound to them using Sequel Binding kit 3.0. The prepared SMRTbell template libraries were then sequenced on a PacBio Sequel sequencer using v3 sequencing primer, 1M v3 SMRT cells, and v3 sequencing chemistry with 6-hour sequencing movie run times. Filtered for artifacts subread data were assembled with Falcon version pb-assembly=0.0.2|falcon-kit=1.2.3|pypeflow=2.1.0 (https://github.com/PacificBiosciences/FALCON) and polished with Arrow version SMRTLink v7.0.1.66975 (https://www.pacb.com/support/software-downloads).

For the genomes of *Leucogyrophana mollusca* and *Hygrophoropsis aurantiaca*, 100 ng of DNA was sheared to 300 bp using the Covaris LE220 and size selected using SPRI beads (Beckman Coulter). The fragments were treated with end-repair, A-tailing, and ligation of Illumina compatible adapters (IDT, Inc) using the KAPA-Illumina library creation kit (KAPA biosystems). Genomic reads were first assembled together with Velvet 1.2.07 (Zerbino & Birney, 2008) to create a long mate-pair library with insert 3000 ± 300 bp which was then assembled together with the original Illumina library with AllPathsLG release version R46652 (Gnerre *et al.*, 2011).

For the production of RNA sequences used for gene annotation, stranded cDNA libraries were generated using the Illumina Truseq stranded mRNA Library Prep kit. mRNA was purified from 1 µg of total RNA using magnetic beads containing poly-T oligos. mRNA was fragmented and reversed transcribed using random hexamers and SSII (Invitrogen) followed by second strand synthesis. The fragmented cDNA was treated with end-pair, A-tailing, adapter ligation, and 18-0 cycles of PCR. The prepared libraries were quantified using KAPA Biosystem’s next-generation sequencing library qPCR kit and run on a Roche LightCycler 480 real-time PCR instrument. The library was then sequenced on the Illumina HiSeq sequencing platform utilizing a TruSeq paired-end cluster kit, v4, and Illumina’s cBot instrument to generate a clustered flow cell for sequencing. Sequencing of the flow cell was performed on the Illumina HiSeq 2500 sequencer using HiSeq TruSeq SBS sequencing kits, v4, following a 2x150 indexed run recipe. Sequencing data were then filtered for artifact/process contamination. Transcriptome reads were assembled into consensus sequences using Rnnotator v. 3.3.2 (Martin *et al.*, 2010).

We performed a stringent quality control to avoid any spurious contaminating sequences in the genome assemblies. Any mitochondrial or non-target contaminant contigs were filtered from the final assembly before gene annotation. Contaminants were identified using the following post-assembly methods: BLAST against NCBI databases (nt, refseq.fungi, ref_prok_rep, mitochondrial, NCBI gcontam and Pacbio 2/4K-control sequences), principal component analysis of tetramer nucleotide frequency, contig length x GC x aligned read coverage, alignment of the sample rDNA ITS vs assembled rDNA ITS sequence, BLAST of the assembled rDNA ITS sequences against the UNITE database (https://unite.ut.ee ). Assessments of genome completeness was done by using the CEGMA (Core Eukaryotic Genes Mapping Approach) pipeline and sample EST assembly capture (see Methods S1 for a detailed description). The completeness of the draft assemblies was evaluated using BUSCO v.3.0.2 with default parameters with fungi odb9 gene set (http://buscodev.ezlab.org/datasets/fungiodb9.tar.gz; Simão et al., 2015). Finally, completeness of the genomes was ascertained by comparing the number of PFAM domains from enzymes involved in the primary metabolism (e.g., glycolysis).

All genomes were annotated using the JGI Annotation Pipeline and made available via the JGI fungal genome portal MycoCosm (Grigoriev *et al.*, 2014).

The genomes of *Boletus reticuloceps*, *Butyriboletus roseoflavus*, *Chiua virens* and *Lanmaoa asiatica* were sequenced using the PacBio Sequel platform at GrandOmics Biosciences (Wuhan, China). Fresh fruiting bodies were collected and only pileus (without the hymenophore) and inner stipe tissues were ground in liquid nitrogen. The CTAB method (Doyle & Doyle, 1987) was used for DNA extraction. Two kinds of libraries were constructed before sequencing. The long-read library of 20 kb was prepared using a SMRTbell DNA Template Prep Kit 1.0 (PacBio p/n 10Tal-259-100). The library was purified by the BluePippin (Sage Science, USA). The final library was sequenced on the PacBio Sequel platform (Pacific Biosciences) as described above. In addition, a library of short length inserts (400 bp) was constructed by Illumina TruSeq Nano DNA Library Prep Kits. Four µg genomic DNA was fragmented, then run on a gel and size selected for a range of 300 to 600 bp. The ends of selected DNA fragments were blunted with an A-base overhang and ligated to sequencing adapters. After quality control, the PCR-free library was sequenced on an Illumina HiSeq 2000 sequencing platform with a paired-end sequencing strategy. De novo assembly of filtered PacBio long reads were assembled into contigs with the program Falcon v1.8.1 (Chin *et al.*, 2016). To further improve the accuracy of assembled contigs, two-step polishing strategies were performed: we first used PacBio long reads and carried out an initial polishing with polished with Arrow version SMRTLink v4.0 for one time and then used highly accurate Illumina paired-end reads to further correct the assembly with Pilon v1.22 (Walker *et al.*, 2014) for four times. For the transcriptomes, stranded cDNA libraries were generated using the Illumina Stranded Total RNA Prep kit, ligation with Ribo-Zero Plus and prepared for sequencing on the Illumina HiSeq sequencing platform. All genomes were annotated using the annotation pipeline of GrandOmics Biosciences (Wuhan, China).

Owing to the fact that several genomes were sequenced from genomic DNA extracted from fruiting bodies collected in the field, we performed a stringent quality control to avoid any spurious contaminating sequences in the genome assemblies using the following post-assembly methods: (1) BLASTN query against the NCBI *nt* database; (2) GC-depth analysis was conducted with minimap2 v2.17 (Li, 2018); (3) Alignment of sample rDNA ITS vs assembled rDNA ITS and BLASTN of assembled rDNA ITS against the UNITE database; (4) Evaluation of genome completeness by using CEGMA, mapping of RNA sequence against the genome assembly, and evaluation of sample EST capture by genome assembly.

**Comparing gene repertoires**

Summary statistics for genome assemblies (e.g., genome assembly size, number of contigs and scaffolds, scaffold N50, scaffold L50, number of genes, exons and introns, protein length) are available for each genome at the JGI MycoCosm Boletales portal (<https://mycocosm.jgi.doe.gov/boletales/boletales.info.html>). Gene prediction and functional annotations are also available at this portal including Gene Ontology (GO), Eukaryotic Orthologous Groups of Proteins (KOG), Kyoto Encyclopedia of Genes and Genomes (KEGG), proteases (MEROPS database), and CAZymes. Carbohydrate-active enzymes (CAZymes) were identified using the CAZy database (www.cazy. org) annotation pipeline (Lombard *et al*., 2014) followed by manual curation by the CAZY team. Secreted proteins were identified using our custom pipeline (Pellegrin *et al.,* 2015). Statistics for secretomes and genomic features were calculated and visualized with a custom script, Proteomic Information Navigated Genomic Outlook (*PRINGO*; Miyauchi *et al*. 2020b). We calculated the percentage of variances in genomic features explained by fungal ecology or phylogenetic distances using Permutational multivariate analysis of variance (PERMANOVA) (see Miyauchi *et al.* (2020b) for details). Secondary metabolite clusters (SMCs) were predicted using antiSMASH 5.0 using a relaxed strictness through the online dedicated server (Blin *et al*., 2019). Filtered gene models were used as feature annotations. Resulting .gbz files were analyzed through the BiGSCAPE pipeline using default parameters and the Pfam-A v30.0 database.

Repeated sequences were identified in unmasked genome assemblies downloaded from JGI MycoCosm (Grigoriev *et al.*, 2014) and Nextomics Biosciences (Wuhan, China). Identification and visualization of TEs were conducted by using a custom workflow script, Transposon Identification Nominative Genome Overview (*TINGO;* Morin *et al.* 2019). For the divergence analyses of TE copies, only high-quality genomes were selected (i.e., the first 10 larger scaffolds should include > 25% of the whole genome assembly). We identified repeat elements with RepeatModeler2 (Flynn *et al*., 2020), which uses three *de novo* repeat finding programs (RECON, RepeatScout and LtrHarvest/Ltr_retriever), followed by annotating the genomes with RepeatMasker (Flynn *et al*., 2020). The copy-divergence analysis was performed using their Kimura 2-parameter distances (K2P) on the selected genomes. K2P distances between genome copies and the consensus models from the library were calculated and plotted using *calcDivergenceFromAlign.pl* and *createRepeatLandscape.pl* (in RepeatMasker util directory), respectively.

**Gene conservation and innovation in Boletales**

To assess the orthology between gene sets from the 28 Boletales species, we downloaded gene models from the JGI MycoCosm database and Nextomics Biosciences (Wuhan, China). We clustered the predicted proteins of these taxa with FASTORTHO using 50% identity, 50% coverage and inflation 3.0 (Wattam *et al.*, 2014). We discussed the protein families (orthogroups) in expansion/contraction in each species relative to the other species only when the differences were statistically supported (Wattam *et al.*, 2014). Based on this clustering, we determined the set of predicted proteins shared by the 28 species (i.e. core genes), sets of predicted proteins encoded in at least two genomes (i.e. dispensable genes) and sets of predicted proteins unique to a genome (i.e. species-specific genes, which are also referred to as taxonomically restricted genes).

Lifestyle-specific genes were manually identified and visualized by using Microsoft Excel. We defined the ectomycorrhiza-specific orthogroups as being present in at least one ectomycorrhizal species of Boletales, but lacking from the genomes of all brown-rot species of Boletales. On the other hand, the brown-rot-specific orthogroups were only found in at least one brown-rot species, but lacking from ectomycorrhizal genomes.

**RNA sequencing of *Phlebopus portentosus***

The strain PP33 of *P. portentosus*, whose genome was sequenced by (Cao *et al.*, 2015), was grown on modified potato dextrose agar (PDA) medium. The culture was then used to inoculate a liquid medium to produce vegetative mycelium for the inoculation on the mixed sawdust/rice seed organic medium (Ji *et al.*, 2011). Total RNA from mycelium on different media were extracted using the TRIzol™ Plus RNA Purification Kit of ThermoFisher. RNA sequencing was performed at Beijing Genomics Institute (BGI) using Illumina HiSeq2000. Raw reads were trimmed and aligned to the reference protein sequences of *P. portentosus* using CLC Genomics Workbench v11. For mapping, the minimum length fraction was 0.9, the minimum similarity fraction 0.9 and the maximum number of hits for a read was set to 10. The unique and total mapped reads number for each transcript were determined and then normalized to RPKM (Reads Per Kilobase of exon model per Million mapped reads). Intact pairs were counted as one, broken pairs were ignored. The normalization of the mapped reads and the differential expression analysis were also conducted with DESeq2 (Love *et al.*, 2014).
